# Estimating temporally variable selection intensity from ancient DNA data with the flexibility of modelling linkage and epistasis

**DOI:** 10.1101/2022.08.02.502360

**Authors:** Zhangyi He, Xiaoyang Dai, Wenyang Lyu, Mark Beaumont, Feng Yu

## Abstract

Innovations in ancient DNA (aDNA) preparation and sequencing technologies have exponentially increased the quality and quantity of aDNA data extracted from ancient biological materials. The additional temporal component from the incoming aDNA data can provide improved power to address fundamental evolutionary questions like characterising selection processes that shape the phenotypes and genotypes of contemporary populations or species. However, utilising aDNA to study past selection processes still involves considerable hurdles like how to eliminate the confounding factor of genetic interactions in the inference of selection. To address this issue, we extend the approach of He et al. (2022) to infer temporally variable selection from the aDNA data in the form of genotype likelihoods with the flexibility of modelling linkage and epistasis in this work. Our posterior computation is carried out by a robust adaptive version of the particle marginal Metropolis-Hastings algorithm with a coerced acceptance rate. Our extension inherits the desirable features of He et al. (2022) such as modelling sample uncertainty resulting from the damage and fragmentation of aDNA molecules and reconstructing underlying gamete frequency trajectories of the population. We evaluate its performance through extensive simulations and show its utility with an application to the aDNA data from pigmentation loci in horses.

## 1. Introduction

Natural selection is one of the primary mechanisms of evolutionary changes and is responsible for the evolution of adaptive features (Darwin, 1859). A full understanding of the role of selection in driving evolutionary changes needs accurate estimates of the underlying timing and strength of selection. With recent advances in sequencing technologies and molecular techniques tailored to ultra-damaged templates, high-quality time serial samples of segregating alleles have become increasingly common in ancestral populations, (*e.g*., Mathieson et al., 2015; Loog et al., 2017; Fages et al., 2019; Alves et al., 2019). The additional temporal dimension of the ancient DNA (aDNA) data has the promise of boosting power of estimating population genetic parameters, in particular for the pace of adaptation, as the allele frequency trajectory through time itself gives us valuable information collected before, during and after genetic changes driven by selection. See Dehasque et al. (2020) for a review of the inference of selection from aDNA.

The temporal component provided by the incoming aDNA data spurred the development of statistical approaches for the inference of selection from time series data of allele frequencies in the last 15 years (see Malaspinas, 2016, for a review). Most existing methods are built upon the hidden Markov model (HMM) framework of Williamson & Slatkin (1999), where the population allele frequency is modelled as a hidden state evolving under the Wright-Fisher model (Fisher, 1922; Wright, 1931), and the sample allele frequency drawn from the underlying population at each given time point is modelled as a noisy observation of the population allele frequency (see Tataru et al., 2017, for a review of statistical inference in the Wright-Fisher model based on time series data of allele frequencies). However, such an HMM framework can be computationally infeasible for large population sizes and evolutionary timescales owing to a prohibitively large amount of computation and storage required in its likelihood calculations.

To our knowledge, most existing methods tailored to aDNA are based on the diffusion limit of the Wright-Fisher model. By working with the diffusion approximation, their HMM framework permits efficient integration over the probability distribution of the underlying population allele frequencies, and therefore, the calculation of the likelihood based on the observed sample allele frequencies can be completed within a reasonable amount of time (e.g., Bollback et al., 2008; Malaspinas et al., 2012; Steinrücken et al., 2014; Schraiber et al., 2016; Ferrer-Admetlla et al., 2016; He et al., 2020b,c, 2022; Lyu et al., 2022). These methods have been successfully applied in aDNA studies, *e.g*., Ludwig et al. (2009) used the method of Bollback et al. (2008) to analyse the aDNA data from pigmentation loci in horses and found that positive selection acted on the derived *ASIP* and *MC1R* alleles, showing that domestication and selective breeding contributed to changes in horse coat colouration.

Despite the availability of a certain number of statistical methods for the inference of selection from genetic time series, their application to aDNA data from natural populations remains limited. Most existing methods are developed in the absence of genetic interactions like linkage and epistasis, with the exception of He et al. (2020b). Linkage and recombination are explicitly modelled in He et al. (2020b), which has been illustrated to significantly improve the inference of selection, in particular for tightly linked loci. Ignoring epistasis can also cause severe issues in the study of selection since the combined effects of mutant alleles may be impossible to predict according to the measured individual effects of a given mutant allele (Bank et al., 2014). As an example, the base coat colours of horses are determined by *ASIP* and *MC1R*. The derived *ASIP* and *MC1R* alleles have been shown to be positively selected through existing approaches (e.g., Bollback et al., 2008; Malaspinas et al., 2012; Steinrücken et al., 2014; Schraiber et al., 2016; He et al., 2020c). However, this is not sufficient to conclude that black horses are favoured by selection since the alleles at *MC1R* interact epistatically with those at *ASIP, i.e*., the presence of at least one copy of the dominant ancestral allele at *MC1R*, and the resulting production of black pigment, is required to check the action of the alleles at *ASIP* (Corbin et al., 2020).

To circumvent this issue, in this work we introduce a novel Bayesian method for the inference of selection acting on the phenotypic trait, allowing the intensity to vary over time, from data on aDNA sequences, with the flexibility of modelling genetic linkage and epistatic interaction. Our method is built upon the two-layer HMM framework of He et al. (2022), and our key innovation is to introduce a Wright-Fisher diffusion that can model the dynamics of two linked genes under phenotypic selection over time to be the underlying Markov process, which permits linkage and epistasis. To remain computationally feasible, our posterior computation is carried out with the particle marginal Metropolis-Hastings (PMMH) algorithm developed by Andrieu et al. (2010), where we adopt the adaptation strategy introduced by Vihola (2012) to tune the covariance structure of the proposal to achieve a given acceptance rate. Also, our approach inherits certain desirable features from He et al. (2022) such as modelling sample uncertainty due to the damage and fragmentation of aDNA molecules and reconstructing underlying frequency trajectories of the gametes in the population. We use our method to analyse the aDNA data associated with horse base coat colours (determined by *ASIP* and *MC1R* with epistatic interaction) and horse pinto coat patterns (controlled by *KIT13* and *KIT16* with tight linkage) from Wutke et al. (2016) to show it applicability, and its performance is evaluated with various simulated data.

## 2. Materials and Methods

In this section, we construct a Wright-Fisher model to characterise two linked genes evolving under phenotypic selection over time first and then derive its diffusion limit. Working with the diffusion approximation, we extend the approach of He et al. (2022) to infer temporally variable selection from the data on aDNA sequences while modelling linkage and epistasis.

### 2.1. Wright-Fisher diffusion

We consider a population of randomly mating diploid individuals represented by alleles at loci 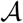 and 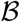 evolving under selection with discrete non-overlapping generations. At each locus, there are two possible allele types, labelled 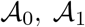 and 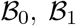, respectively, resulting in four possible haplotypes on both loci, 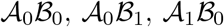 and 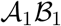, labelled haplotypes 00, 01, 10 and 11, respectively. We attach the symbols 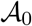 and 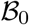 to the ancestral alleles, which we assume originally exist in the population, and we attach the symbols 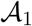 and 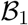 to the mutant alleles, which we assume arise only once in the population. Considering the absence of sex effects, this setup gives rise to 10 possible (unordered) genotypes 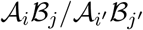, corresponding to at most 10 distinct phenotypes 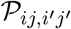. Phenotypes 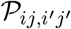 and 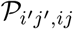 are identical in our notation.

We incorporate viability selection into the population dynamics and assume that the viability is only determined by the phenotype. Viabilities of all phenotypes at loci 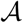 and 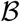 per generation are assigned 1 + *s_ij,i′j′_*, where *s_ij,i′j′_* is the selection coefficient of the 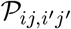 phenotype with *s_ij,i′j′_* ∈ [−1, +∞) and *s_ij,i′j′_* = *s_i′j′,ij_*. In what follows, we let the selection coefficient *s*_00,00_ = 0 unless otherwise noted, and then *s_ij,i′j′_* denotes the selection coefficient of the 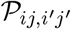 phenotype against the 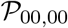 phenotype.

#### 2.1.1. Wright-Fisher model

Let 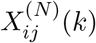 denote the gamete frequency of haplotype *ij* at generation 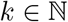 and ***X***^(*N*)^(*k*) be the vector of the four gamete frequencies. To incorporate non-constant demographic histories, we assume that the population size changes deterministically, with *N*(*k*) denoting the number of diploid individuals in the population at generation *k*. In the Wright-Fisher model, we assume that gametes are randomly chosen from an effectively infinite gamete pool reflecting the parental gamete frequencies at each generation. We therefore have

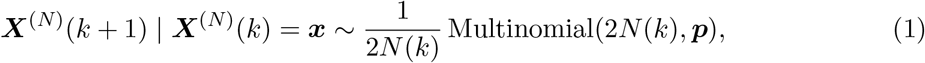

where ***p*** is the vector of parental gamete frequencies. Under the assumption of random mating, we can further express the vector of parental gamete frequencies as

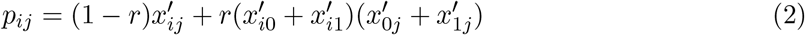

for *i, j* ∈ {0, 1}, where

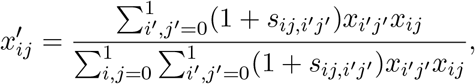

and *r* denotes the recombination rate of the 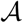 and 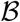 loci located on the same chromosome, *i.e*., the fraction of recombinant offspring showing a crossover between the two loci per generation. If the 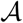 and 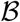 loci are located on separate chromosomes, we let the (artificial) recombination rate *r* = 0.5 (i.e., free recombination). The two-locus Wright-Fisher model with selection is defined as the Markov process ***X***^(*N*)^ evolving with transition probabilities in Eq. (1) in the state space 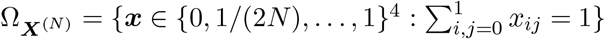.

#### 2.1.2. Diffusion approximation

We study the two-locus Wright-Fisher model with selection through its diffusion limit due to the complicated nature of its transition probability matrix, in particular for large population sizes or evolutionary timescales. More specifically, we measure time in a unit of 2N0 generations, denoted by *t*, where *N*_0_ is an arbitrary reference population size fixed through time, and assume that the selection coefficients and recombination rate are all of order 1/(2*N*_0_). As the reference population size *N*_0_ approaches infinity, the scaled selection coefficients *α_ij,i′j′_* = 2*N*_0_*s_ij,i′j′_* and the scaled recombination rate *ρ* = 4*N*_0_*r* are kept constant, and the ratio of the population size to the reference population size *N*(*t*)/*N*_0_ converges to a function, denoted by *β*(*t*). Notice that the assumption will be violated if the 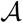 and 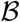 loci are located on separate chromosomes, *i.e*., *r* = 0.5, but we shall nevertheless use this scaling to find the drift term in the diffusion limit. We will plug the unscaled recombination rate *r* into the resulting system of stochastic differential equations (SDE’s) and use that as our diffusion approximation.

Let 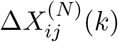 denote the change in the gamete frequency of haplotype *ij* over generation *k*. With standard techniques of diffusion theory (see, *e.g*., Karlin & Taylor, 1981), we can formulate the infinitesimal mean vector ***μ***(*t, **x***) and the infinitesimal (co)variance matrix **Σ**(*t, **x***) as

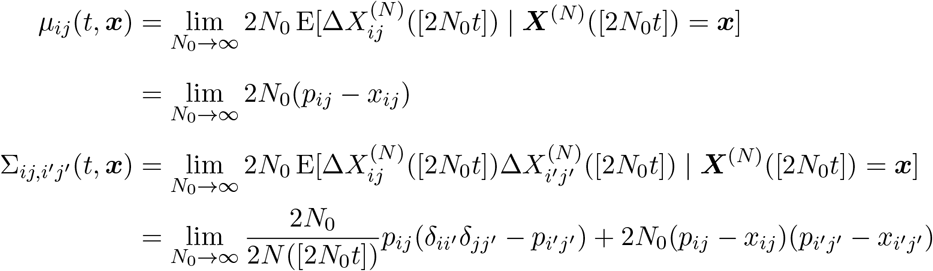

for *i, j, i′, j′* ∈ {0, 1}, where *δ* denotes the Kronecker delta function and [·] is used to represent the integer part of the value in the brackets.

To obtain the expression for the infinitesimal mean vector ***μ***(*t, **x***), we compute the limit of the expected change in the gamete frequency of haplotype *ij* within a single generation as the reference population size *N*_0_ goes to infinity. The only terms that survive after taking the limit are the first order terms in the Taylor expansion of the sampling probability *p_ij_* in Eq. (2) with respect to the selection coefficients *s_ij,i′j′_* and recombination rate *r*. We have

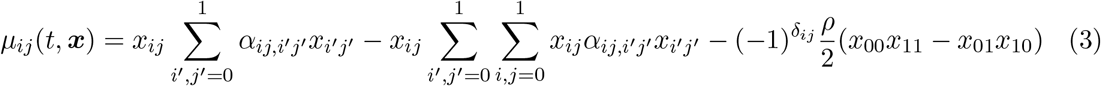

for *i, j* ∈ {0, 1}. Note that we take the scaled recombination rate to be *ρ* = 2*N*_0_ (*i.e*., the (artificial) recombination rate *r* = 0.5) if the 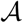 and 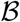 loci are located on separate chromosomes. Such a strong recombination term serves to uncouple the two genes located on separate chromosomes. The infinitesimal (co)variance matrix **Σ**(*t, **x***) corresponds to the standard Wright-Fisher diffusion on four haplotypes (see, e.g., He et al., 2020a). We have

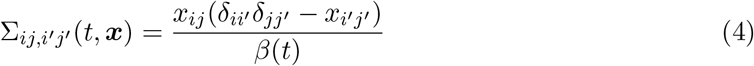

*i, j, i′, j′* ∈ {0, 1}.

Combining the Wright-Fisher diffusion with the infinitesimal mean vector ***μ***(*t, **x***) in Eq. (3) and the infinitesimal (co)variance matrix **Σ**(*t, **x***) in Eq. (4), we achieve the following system of SDE’s as our diffusion approximation of the Wright-Fisher model in Eq. (1)

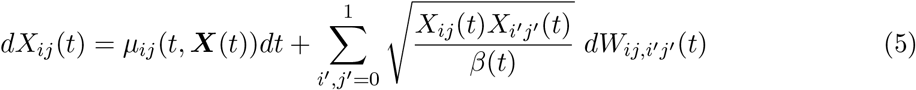

for *i, j* ∈ {0, 1}, where *W_ij,i′j′_* denotes an independent standard Wiener process with *W_ij,i′j′_*(*t*) = –*W_i′j′,ij_*(*t*). This anti-symmetry requirement implies *W_ij,ij_*(*t*) = 0, and the (co)variance matrix for the *X_ij_*’s is exactly the infinitesimal (co)variance matrix **Σ**(*t, **x***) in Eq. (4). We refer to the diffusion process ***X*** evolving in the state space 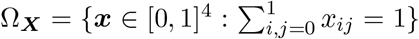 that solves the system of SDE’s in Eq. (5) as the two-locus Wright-Fisher diffusion with selection.

### 2.2. Bayesian inference of selection

Suppose that the available data are always sampled from the underlying population at a finite number of distinct time points, say *t*_1_ < *t*_2_ < … < *t_K_*, measured in units of 2*N*_0_ generations. We assume that *N_k_* individuals are drawn from the underlying population at the *k*-th sampling time point, and for individual *n*, let ***r**_l,n,k_* be, in this generic notation, all of the reads at locus *l* for *l* ∈ {1, 2}. The population genetic quantities of our interest are the selection coefficients *s_ij,i′j′_* for *i, j, i′, j′* ∈ {0, 1}. Recall that our setup gives rise to at most 10 distinct phenotypes (i.e., at most 9 distinct selection coefficients). For simplicity, we use to represent all distinct selection coefficients to estimate.

#### 2.2.1. Hidden Markov model

We extend the two-layer HMM framework of He et al. (2022) to model linkage and epistasis, where the first hidden layer ***X***(*t*) characterises the gamete frequency trajectories of the underlying population through time by the Wright-Fisher diffusion in Eq. (5), the second hidden layer ***G***(*t*) represents the genotype of the individual in the sample, and the third observed layer ***R***(*t*) denotes the data on ancient DNA sequences (see Figure 1).

**Figure 1:**
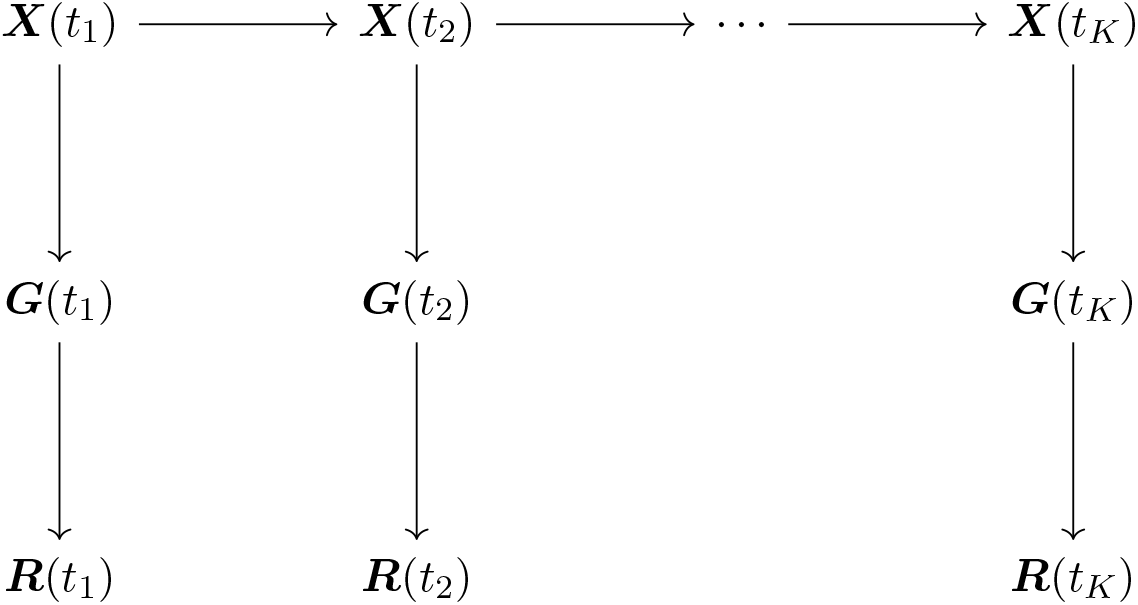
Graphical representation of the two-layer HMM framework extended from He et al. (2022) for the data on ancient DNA sequences.

We let ***x***_1:*K*_ = {*x*_1_, *x*_2_,…, *x_K_*} be the frequency trajectories of the gametes in the underlying population at the sampling time points ***t***_1:*K*_ and ***g***_1:*K*_ = {***g***_1_, ***g***_2_,…, ***g**_K_*} be the genotypes of the individuals drawn from the underlying population at the sampling time points ***t***_1:*K*_, where ***g**_k_* = {***g***_1,*k*_, ***g***_2,*k*_,…, ***g**_N_k__,k*} with ***g**_n,k_* = {*g*_1,*n,k*_, *g*_2,*n,k*_} and *g_l,n,k_* ∈ {0, 1, 2} being the number of mutant alleles at locus *l* in individual *n* at sampling time point *t_k_*. Based on the HMM framework illustrated in Figure 1, the posterior probability distribution for the selection coefficients and population gamete frequency trajectories can be expressed as

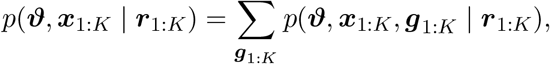

where

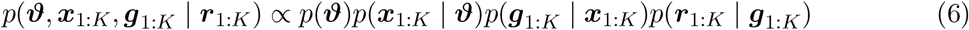

and ***r***_1:*K*_ = {***r***_1_, ***r***_2_,…, *r_K_*} with *r_k_* = {***r***_1,*k*_, ***r***_2,*k*_,…, ***r**_N_k__,k*} and *r_n,k_* = {***r***_1,*n,k*_, ***r***_2,*n,k*_}.

The first term of the product in Eq. (6), 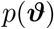 is the prior probability distribution for the selection coefficients. We can adopt a uniform prior over the interval [−1, +∞) for each selection coefficient if our prior knowledge is poor.

The second term of the product in Eq. (6), 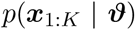, is the probability distribution for the population gamete frequency trajectories at all sampling time points. As the Wright-Fisher diffusion is a Markov process, we can decompose the probability distribution 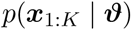 as

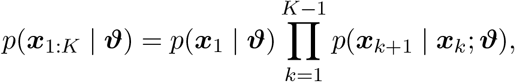

where 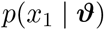 is the prior probability distribution for the population gamete frequencies at the initial sampling time point, set to be a flat Dirichlet distribution over the state space Ω***_X_*** if our prior knowledge is poor, and 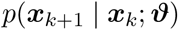 is the transition probability density function of the Wright-Fisher diffusion X between two consecutive sampling time points for *k* = 1, 2,…, *K* – 1, solving the Kolmogorov backward equation (or its adjoint) associated with the Wright-Fisher diffusion in Eq. (5).

The third term of the product in Eq. (6), *p*(***g***_1:*K*_| ***x***_1:*K*_), is the probability distribution for the genotypes of all individuals in the sample given the population gamete frequency trajectories at all sampling time points. With the conditional independence from our HMM framework (see Figure 1), we can decompose the probability distribution *p*(***g***_1:*K*_| ***x***_1:*K*_) as

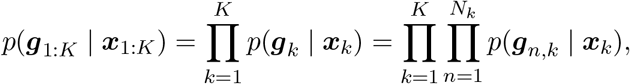

where *p*(***g**_n,k_*| ***x**_k_*) is the probability distribution for the genotypes ***g**_n,k_* of sampled individual *n* given the gamete frequencies ***x**_k_* of the population. Under the assumption that all individuals in the sample are drawn from the population in their adulthood (*i.e*., the stage after selection but before recombination in the life cycle, see He et al. (2017)), the probability of observing the sampled individual genotypes ***g**_n,k_* = (*i + i′, j + j′*) given the population gamete frequencies can be calculated with

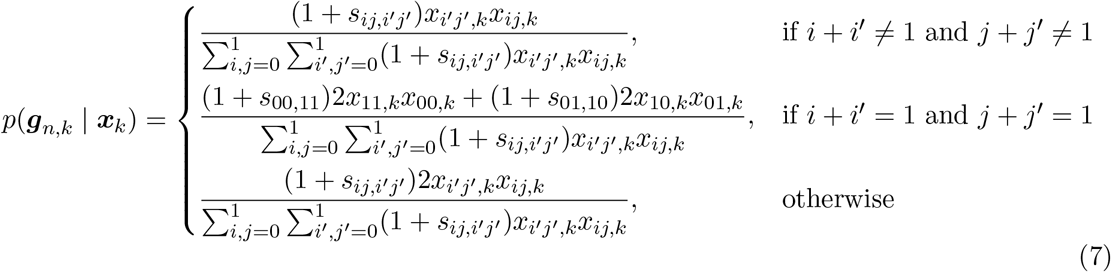

for *i, j, i′ j′* = 0, 1.

The fourth term of the product in Eq. (6), *p*(***r***_1:*K*_ | ***g***_1:*K*_), is the probability of observing the reads of all sampled individuals given their corresponding genotypes. Using the conditional independence from our HMM framework, as shown in Figure 1, we can decompose the probability *p*(***r***_1:*K*_ | ***g***_1:*K*_) as

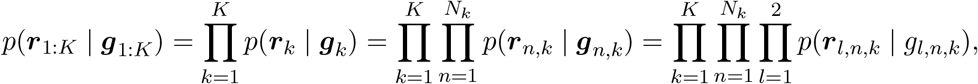

where *p*(***r**_l,n,k_* | ***g**_l,n,k_*) is the probability of observing the reads ***r**_l,n,k_* of sampled individual n at locus l given its genotype *g_l,n,k_*, known as the genotype likelihood, which is commonly available with aDNA data.

#### 2.2.2. Adaptive particle marginal Metropolis-Hastings

Similar to He et al. (2022), we carry out our posterior computation by the PMMH algorithm (Andrieu et al., 2010) that enables us to jointly update the selection coefficients and population gamete frequency trajectories. More specifically, we estimate the marginal likelihood

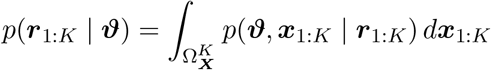

through the bootstrap particle filter (Gordon et al., 1993), where we generate the particles from the Wright-Fisher SDE’s in Eq. (5) by the Euler-Maruyama scheme. The product of the average weights of the set of particles at the sampling time points ***t***_1:*K*_ yields an unbiased estimate of the marginal likelihood 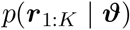, denoted by 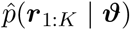. The population gamete frequency trajectories ***x***_1:*K*_ are sampled once from the final set of particles with their relevant weights.

Although the PMMH algorithm has been shown to work well in He et al. (2022), in practice, its performance depends strongly on the choice of the proposal. In this work, due to the increase in the number of selection coefficients required to be estimated, choosing an appropriate proposal to ensure computational efficiency becomes challenging. To resolve this issue, we adopt a random walk proposal with covariance matrix **Γ**, denoted by 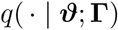, the Gaussian probability density function with mean vector 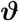 and covariance matrix **Γ**, and under ideal conditions, the optimal choice of the covariance matrix **Γ** is a rescaled version of the covariance matrix of the posterior (Roberts & Rosenthal, 2001). Given that the covariance matrix of the posterior is commonly not available in advance, we adopt the adaptation strategy (Vihola, 2012) that can dynamically align the covariance matrix of the proposal with that of the posterior based on accepted samples. More specifically, we prespecify a target acceptance rate, denoted by *A\**, and a step size sequence (decaying to zero), denoted 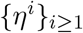, where the superscript denotes the iteration. The covariance matrix is updated by following the iteration formula

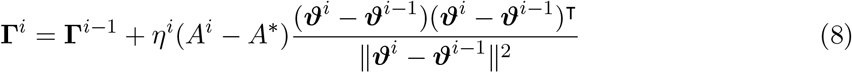

with the covariance matrix **Γ**^1^ (*e.g*., **Γ**^1^ = *σ*^2^***I***) and selection coefficients 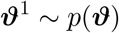, where

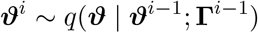

and

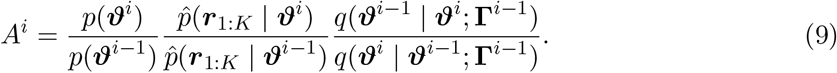

Such an adaptation strategy can also coerce the acceptance rate. In practice, the target acceptance rate is set to *A\** ∈ [0.234, 0.440], and the step size sequence is defined as *η^i^* = *i^-γ^* with *γ* ∈ (0.5, 1] (Vihola, 2012). See Luengo et al. (2020) and references therein for other adaptation strategies.

For the sake of clarity, we write down the robust adaptive version of the PMMH algorithm for our posterior computation:

Step 1: Initialise the selection coefficients 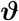 and population gamete frequency trajectories ***x***_1:*K*_:

Step 1a: Draw 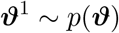.
Step 1b: Run a bootstrap particle filter with 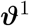 to get 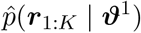 and 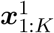.
Step 1c: Initialise **Γ**^i^. Repeat Step 2 until enough samples of the selection coefficients 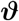 and population gamete frequency trajectories ***x***_1:*K*_ have been attained:
Step 2: Update the selection coefficients 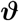 and population gamete frequency trajectories ***x***_1:*K*_:

Step 2a: Draw 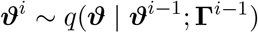.
Step 2b: Run a bootstrap particle filter with 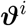 to get 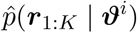 and 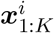.
Step 2c: Update **Γ***^i^* through Eqs. (8) and (9).
Step 2d: Accept 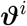 and 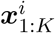 with *A^i^* and set 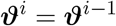 and 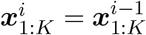 otherwise.

With enough samples of the selection coefficients 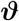 and population gamete frequency trajectories ***x***_1:*K*_, we produce the minimum mean square error estimates for the selection coefficients 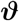 and population gamete frequency trajectories ***x***_1:*K*_ by calculating their posterior means.

As in He et al. (2022), our procedure can allow the selection coefficients *s_ij,i′j′_* to change over time (piecewise constant), e.g., let the selection coefficients 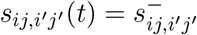 if *t* < *τ* otherwise 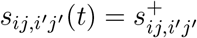, where *τ* is the time of an event that might change selection, e.g., the times of plant and animal domestication. The only modification required is to simulate the population gamete frequency trajectories ***x***_1:*K*_ according to the Wright-Fisher diffusion with the selection coefficients 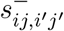 for *t* < *τ* and 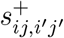 for *t* ≥ *τ*, respectively. In this setup, we propose a scheme to test the hypothesis whether selection changes at time *τ* for each phenotypic trait, including estimating their selection differences, through computing the posterior *p*(Δ*s_ij,i′j′_* | ***r***_1:*K*_) from the PMMH samples of the selection coefficients 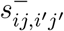 and 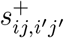, where 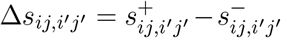, denotes the change in the selection coefficient at time *τ*. Note that our method can handle the case that the events that might change selection are different for different phenotypic traits (*i.e*., the time *τ* could be taken to be different values for different phenotypic traits).

## 3. Results

In this section, we employ our approach to reanalyse the published ancient horse DNA data from earlier studies of Ludwig et al. (2009), Pruvost et al. (2011) and Wutke et al. (2016), where they sequenced 201 ancient horse samples in total ranging from a pre- to a post-domestication period for eight loci coding for horse coat colouration. In particular, we perform the inference of selection that acts on the base coat colour determined by *ASIP* and *MC1R* and the pinto coat pattern controlled by *KIT13* and *KIT16* and test the hypothesis that there was a shift in human preferences for horse coat colours and patterns in the Middle Ages when horses started to be differentiated by use (Wutke et al., 2016). Various simulation studies, assessing the performance of our procedure, are available in the supplement.

It has been shown by Gaunitz et al. (2018) that the ancient horse samples providing the earliest archaeological evidence of domestication (occurred 5500 years ago) are not the ancestors of modern domestic horses, which only spread out from the Pontic–Caspian steppe 4200 years ago (the domestication of modern horses) when they rapidly started to become dominant across Eurasia (Librado et al., 2021). Also, horse populations outside the Pontic–Caspian steppe and older than 4200 years ago show strong geographic structure, e.g., horse populations from Iberia and Siberia are completely different (Fages et al., 2019). Hence, to eliminate these confounding effects, we exclude the ancient horse samples older than 4200 years ago in our analyses, but we assume that the respective mutations occurred at all pigmentation loci 4200 years ago.

As Wutke et al. (2016) only provided called genotypes for each gene (including missing calls), we use the same scheme as in He et al. (2022) to convert to corresponding genotype likelihoods. More specifically, we take the genotype likelihood of the called genotype to be 1 and those of the remaining two to be 0 if the genotype is called, and otherwise, all possible (ordered) genotypes are assigned equal genotype likelihoods (normalised to sum to 1). Genotype likelihoods for each gene can be found in Table S1.

In what follows, we set the average length of a generation of the horse to be eight years and use the time-varying size of the horse population estimated by Der Sarkissian et al. (2015) (see Figure S1) with the reference population size *N*_0_ = 16000 (*i.e*., the most recent population size) like Schraiber et al. (2016) unless otherwise noted. Since the flat Dirichlet prior for the starting population gamete frequencies is more likely to produce low linkage disequilibrium, we generate the starting population gamete frequencies x_1_ through the following procedure:

Step 1: Draw *y*_1_, *y*_2_ ~ Uniform(0, 1).
Step 2: Draw *D* ~ Uniform(max{–*y*_1_*y*_2_, –(1 – *y*_1_)(1 – *y*_2_)}, min{*y*_1_(1 – *y*_2_), (1 – *y*_1_)*y*_2_}).
Step 3: Set ***x***_1_ = ((1 – *y*_1_)(1 – *y*_1_) + D, (1 – *y*_1_)*y*_2_ – *D*, *y*_1_(1 – *y*_2_) – *D*, *y*_1_*y*_2_ + *D*).

Note that *y*_1_ and *y*_2_ denote the starting population frequencies of the mutant allele at the two loci, respectively, and D is the coefficient of linkage disequilibrium. We run our adaptive PMMH algorithm with 1000 particles and 20000 iterations, where we set the target acceptance rate to *A\** = 0.4 and define the step size sequence as *η_i_* = *i*^-2/3^ for i = 1,2,…, 20000. We divide each generation into five subintervals in the Euler-Maruyama scheme. We discard a burn-in of 10000 iterations and thin the remaining iterations by keeping every fifth value.

### 3.1. Horse base coat colours

The horse genes *ASIP* and *MC1R* are primarily responsible for determination of base coat colours (i.e., bay, black and chestnut). *ASIP*, located on chromosome 22, is epistatic to *MC1R*, located on chromosome 3 (Rieder et al., 2001). At each locus, there are two allele types, labelled *A* and *a* for *ASIP* and *E* and *e* for *MC1R*, respectively, where the capital letter represents the ancestral allele and the small letter represents the mutant allele. See Table 1 for the genotype-phenotype map at *ASIP* and *MC1R* for base coat colours.

**Table 1:** The genotype-phenotype map at *ASIP* and *MC1R* for horse base coat colours.

|  |  | *MC1R* |  |  |
| --- | --- | --- | --- | --- |
|  |  | *E/E* | *E/e* | *e/e* |
| *ASIP* | *A/A* | bay | bay | chestnut |
|  | *A/a* | bay | bay | chestnut |
|  | *a/a* | black | black | chestnut |

#### 3.1.1. Wright-Fisher diffusion for ASIP and MC1R

Let us consider a horse population represented by the alleles at *ASIP* and *MC1R* evolving under selection over time, which induces four possible haplotypes *AE*, *Ae*, *aE* and *ae*, labelled haplotypes 00, 01, 01 and 11, respectively. We take the relative viabilities of the three phenotypes, i.e., the bay, black and chestnut coat, to be 1, 1 + *s_b_* and 1 + *s_c_*, respectively, where *s_b_* is the selection coefficient of the black coat against the bay coat and *s_c_* is the selection coefficient of the chestnut coat against the bay coat. See Table 2 for the relative viabilities of all genotypes at *ASIP* and *MC1R*.

**Table 2:** Relative viabilities of all genotypes at *ASIP* and *MC1R*.

|  | *AE* | *Ae* | *aE* | *ae* |
| --- | --- | --- | --- | --- |
| *AE* | 1 | 1 | 1 | 1 |
| *Ae* | 1 | $1 + s_c$ | 1 | $1 + s_c$ |
| *aE* | 1 | 1 | $1 + s_b$ | $1 + s_b$ |
| *ae* | 1 | $1 + s_c$ | $1 + s_b$ | $1 + s_c$ |

We measure time in units of 2*N*_0_ generations and scale the selection coefficients *α_b_* = 2*N*_0_*s_b_, α_c_* = 2*N*_0_*s_c_* and recombination rate *ρ* = 4*N*_0_*r*, respectively. Let *X_ij_*(*t*) be the gamete frequency of haplotype *ij* at time *t*, which satisfies the Wright-Fisher SDE’s in Eq. (5). More specifically, the drift term ***μ***(*t, **x***) can be simplified with the genotype-phenotype map shown in Table 2 as

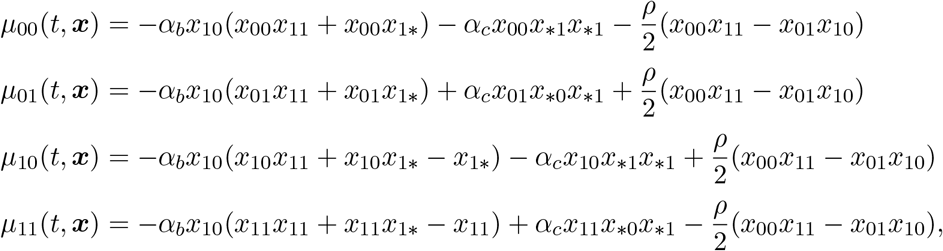

where we take the scaled recombination rate to be *ρ* = 2*N*_0_ since the two genes are located on separate chromosomes.

#### 3.1.2. Selection of horse base coat colours

Given that *ASIP* and *MC1R* are located on separate chromosomes, we generate the starting population gamete frequencies with the procedure described above but fix the coefficient of linkage disequilibrium to zero. The resulting posteriors for the selection coefficients and population phenotype frequency trajectories are shown in Figure 2, and their estimates as well as the 95% highest posterior density (HPD) intervals are summarised in Table S2.

**Figure 2:**
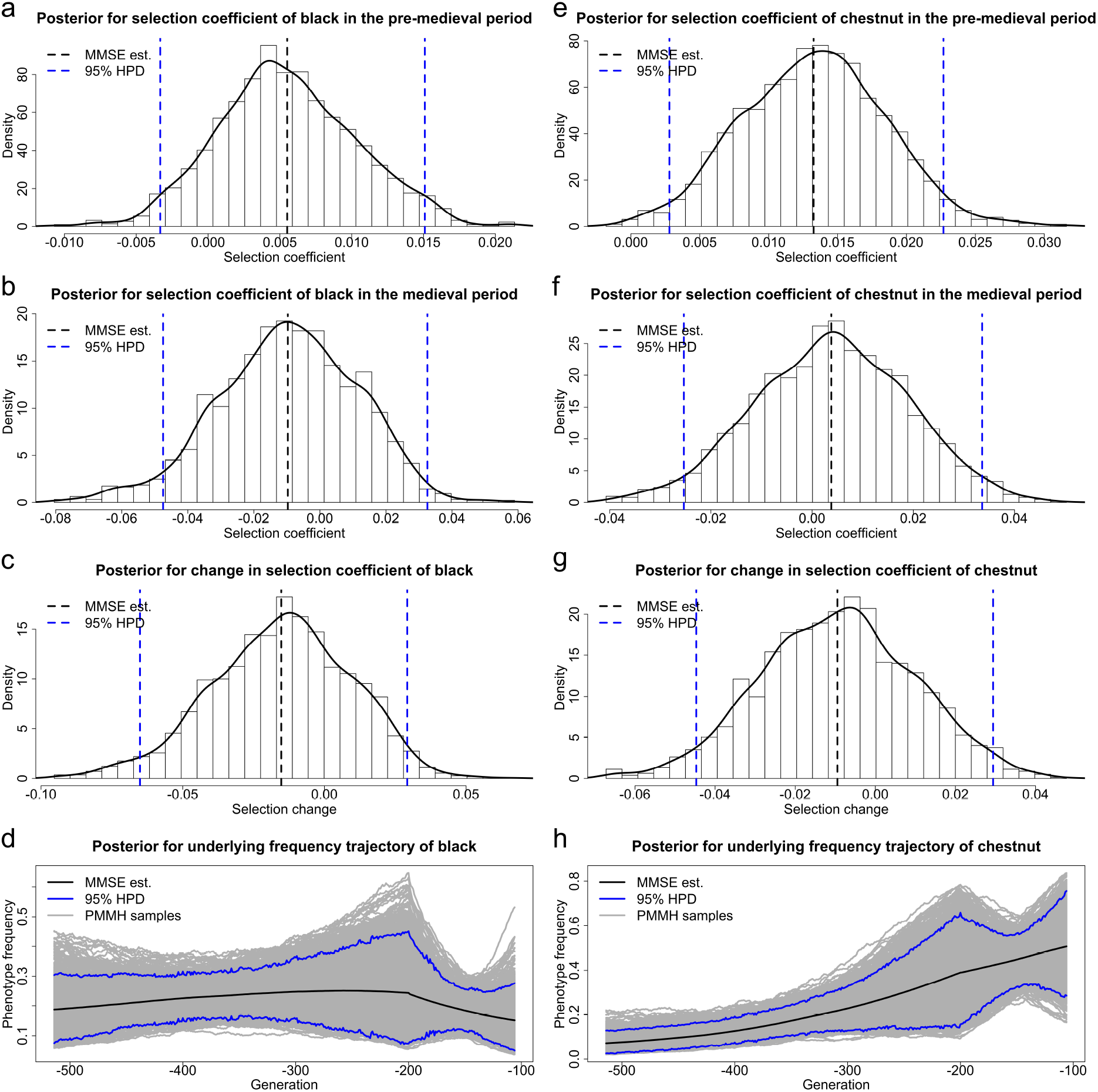
Posteriors for selection of base coat colours in the pre-medieval and medieval period and underlying frequency trajectories of each phenotypic trait in the population, (a)-(d) for the black coat and (e)-(h) for the chestnut coat, respectively.

Our estimate for the selection coefficient 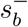 is 0.0055 (95% HPD: [−0.0033, 0.0151]). Though the 95% HPD interval contains 0, we still find that the black coat was most probably favoured by selection in the pre-medieval period (with posterior probability for positive selection being 0.880). Our estimate for the selection coefficient 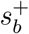 is –0.0097 (95% HPD: [−0.0475, 0.0326]), providing weak evidence that the black coat was selectively deleterious in the medieval period (with posterior probability for negative selection being 0.666). From Figure 2c, we observe that a negative shift took place in selection of the black coat in Middle Ages (with posterior probability for a negative change being 0.728). Our estimate for the underlying frequency trajectory of the black coat reveals a slow increase in the pre-medieval period and then a slight decrease in the medieval period.

Our estimate for the selection coefficient 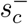 is 0.0133 (95% HPD: [−0.0028, 0.0227]), providing enough evidence to support that the chestnut coat was positively selected in the pre-medieval period (with posterior probability for positive selection being 0.998). Our estimate for the selection coefficient 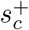 is 0.0038 (95% HPD: [−0.0254, 0.0336]), suggesting that in the medieval period the chestnut coat was still favoured by selection (with posterior probability for positive selection being 0.613). Figure 2g shows that a negative shift occurred in selection of the chestnut coat in Middle Ages (with posterior probability for a negative change being 0.698). We see from Figure 2e that the underlying frequency of the chestnut keeps increasing from the domestication of modern horses.

See Figure S2 and Table S3 for the results produced with a flat Dirichlet prior for the starting population gamete frequencies, which are consistent with those shown in Figure 2.

### 3.2. Horse pinto coat patterns

The horse genes *KIT13* and *KIT16* are mainly responsible for determination of pinto coat patterns (i.e., tobiano and sabino), both of which reside on chromosome 3, 4668 base pairs (bp) apart, with an average rate of recombination 10^-8^ crossover/bp (Dumont & Payseur, 2008). At each locus, there are two allele types, labelled *KM0* for the ancestral allele and *KM1* for the mutant allele at *KIT13* and *sb1* for the ancestral allele and *SB1* for the mutant allele at *KIT16*, respectively. See Table 3 for the genotype-phenotype map at *KIT13* and *KIT16* for pinto coat patterns, where the solid coat pattern refers to a coat that neither tobiano nor sabino is present, and the mixed coat pattern refers to a coat that is a mixture between tobiano and sabino.

**Table 3:**
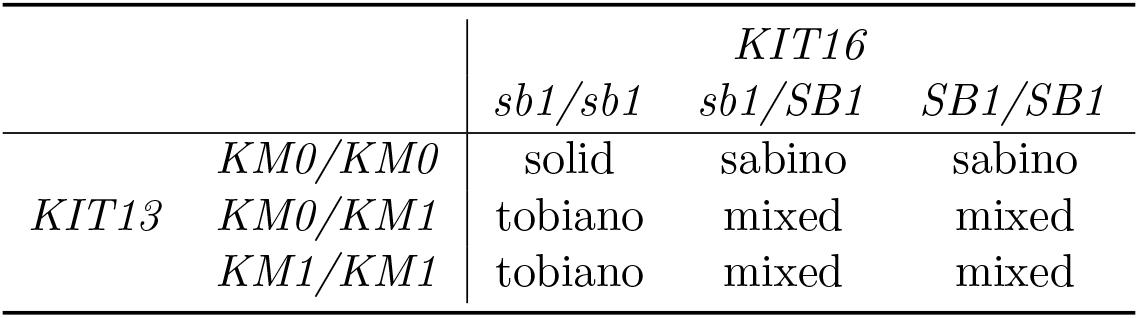
The genotype-phenotype map at *KIT13* and *KIT16* for horse pinto coat patterns.

|  |  | *KIT16* |  |  |
| --- | --- | --- | --- | --- |
|  |  | *sb1/sb1* | *sb1/SB1* | *SB1/SB1* |
| *KIT13* | *KM0/KM0* | solid | sabino | sabino |
|  | *KM0/KM1* | tobiano | mixed | mixed |
|  | *KM1/KM1* | tobiano | mixed | mixed |

#### 3.2.1. Wright-Fisher diffusion for KIT13 and KIT16

We now consider a horse population represented by the alleles at *KIT13* and *KIT16* evolving under selection over time. Such a setup gives rise to four possible haplotypes *KM0sb1, KM0SB1, KM1sb1* and *KM1SB1*, labelled haplotypes 00, 01, 01 and 11, respectively. We take the relative viabilities of the four phenotypes, *i.e*., the solid, tobiano, sabino and mixed coat, to be 1, 1 + *s_to_*, 1 + *s_sb_* and 1 + *s_mx_*, respectively, where *s_to_* is the selection coefficient of the tobiano coat against the solid coat, *s_sb_* is the selection coefficient of the sabino coat against the solid coat, and *s_mx_* is the selection coefficient of the mixed coat against the solid coat. See Table 4 for the relative viabilities of all genotypes at *KIT13* and *KIT16*.

**Table 4:**
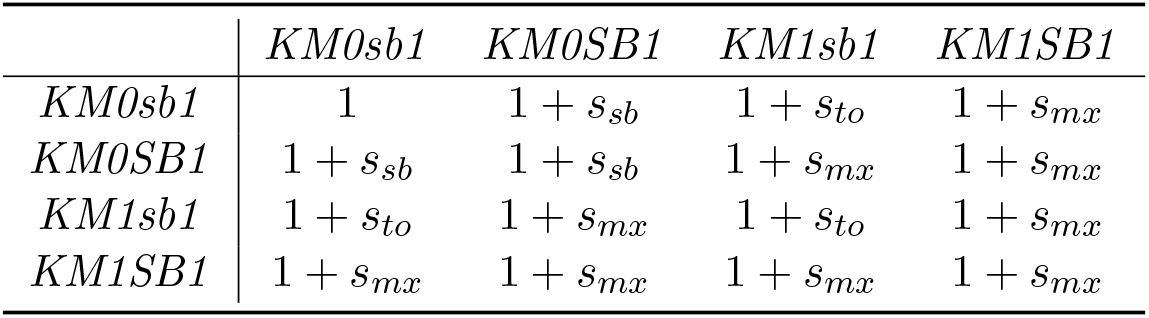
Relative viabilities of all genotypes at *KIT13* and *KIT16*.

We measure time in units of 2*N*_0_ generations and scale the selection coefficients *α_to_* = 2*N*_0_*s_to_, α_sb_* = 2*N*_0_*s_sb_, α_mx_* = 2*N*_0_*s_mx_* and recombination rate *ρ* = 4*N*_0_*r*, respectively. Let *X_ij_* (*t*) be the gamete frequency of haplotype *ij* at time *t*, which follows the Wright-Fisher SDE’s in Eq. (5). In particular, the drift term ***μ***(*t, **x***) can be simplified through Table 4 as

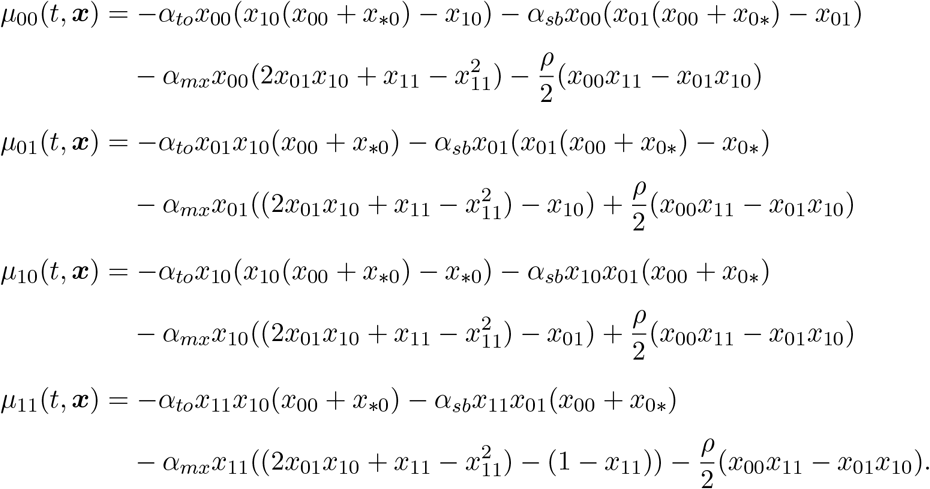

#### 3.2.2. Selection of horse pinto coat patterns

To our knowledge, the mixed coat has never been found in the horse population, so we fix the selection coefficient *s_mx_* = −1 over time. The resulting posteriors for the selection coefficients and population phenotype frequency trajectories are illustrated in Figure 3, and their estimates as well as the 95% HPD intervals are summarised in Table S4.

**Figure 3:**
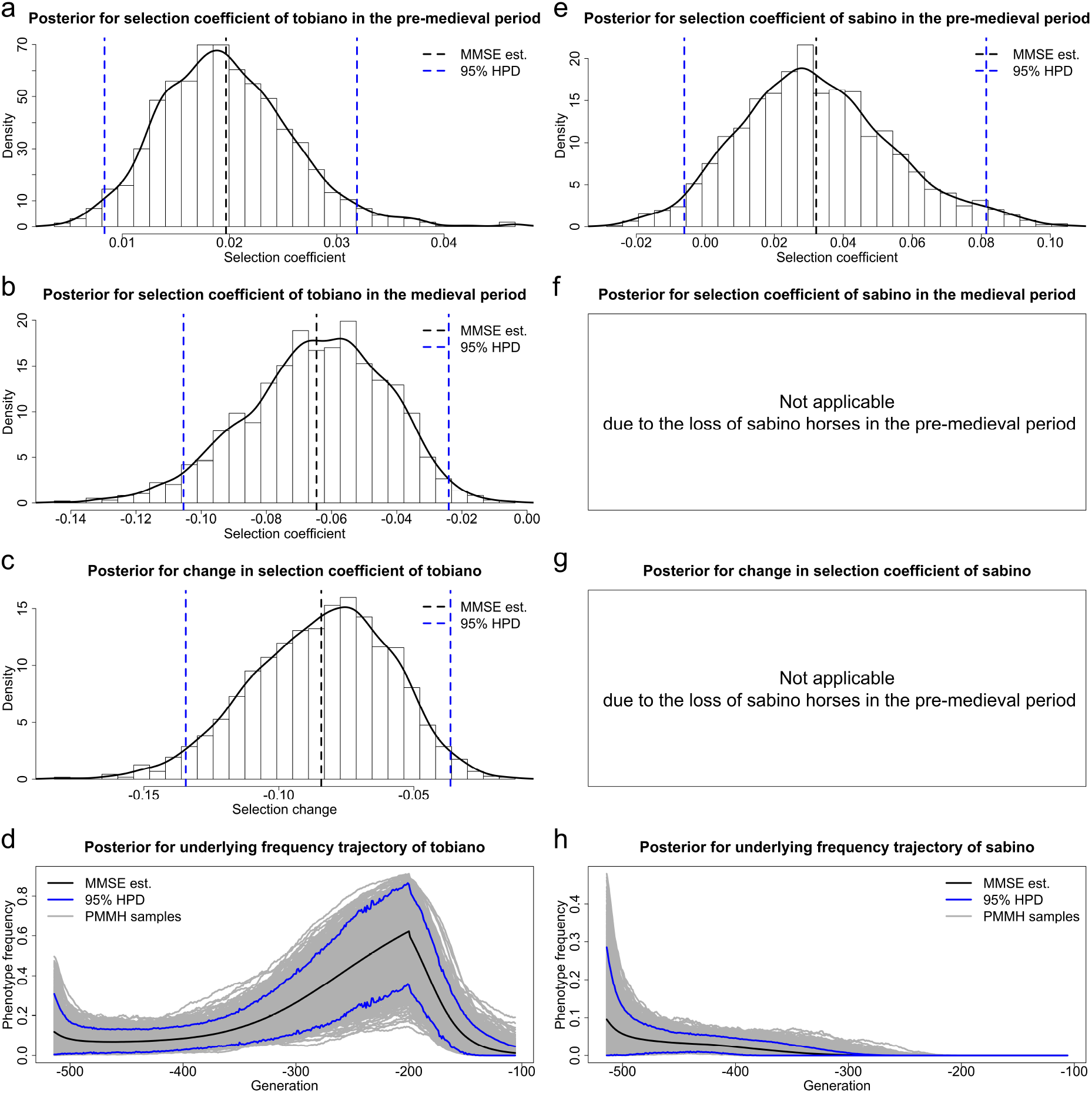
Posteriors for selection of pinto coat patterns in the pre-medieval and medieval period and underlying frequency trajectories of each phenotypic trait in the population, (a)-(d) for the tobiano coat and (e)-(h) for the sabino coat, respectively.

Our estimate for the selection coefficient 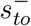 is 0.0197 (95% HPD: [0.0084, 0.0319]) and the selection coefficient 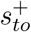 is –0.0646 (95% HPD: [−0.1054, –0.0241]). Our results reveal sufficient evidence to support that the tobiano coat was favoured by selection since the domestication of modern horses (with posterior probability for positive selection being 1) but became selectively deleterious in the medieval period (with posterior probability for negative selection being 1). Figure 3c suggests that a negative shift took place in selection of the tobiano coat in the Middle Ages (with posterior probability for a negative change being 1). Our estimate for the underlying frequency trajectory of the tobiano coat indicates that the frequency of the tobiano coat grows substantially after modern horse domestication and then drops sharply in the medieval period.

Our estimate for the selection coefficient 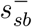 is 0.0321 (95% HPD interval [−0.0061, 0.0814]), which shows compelling evidence of positive selection acting on the sabino coat (with posterior probability for positive selection being 0.941). However, as illustrated in Figure 3c, we observe that the frequency of the sabino coat declines slowly from the domestication of modern horses until the loss of the sabino coat in approximately 24 BC (*i.e*., the earliest time that the upper and lower bounds of the 95% HPD interval for the frequency of the sabino coat are both zero), probably caused by that the sabino coat was somewhat out-competed by the tobiano coat under the tight linkage between *KIT13* and *KIT16*.

Note, we only present the resulting posterior for the selection coefficient 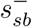. This is because our results show that the sabino coat became extinct in the pre-medieval period (see Figure 3h). Without genetic variation data, our PMMH algorithm fails to converge in reasonable time for the selection coefficient 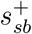, which however has little effect on estimation of the remaining three (see Figure S3, where we repeatedly run our procedure to estimate the selection coefficients 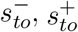 and 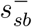 with different prespecified values of the selection coefficient 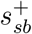, that are uniformly drawn from [−1, 1]).

We also provide the results produced with a flat Dirichlet prior for the starting population gamete frequencies (see Figure S4 and Table S5) and that we co-estimate the selection coefficient *s_mx_* (see Figure S5 and Table S6). Our estimate for the selection coefficient *s_mx_* is −0.5621 (95% HPD: [−0.9645, −0.2262]) in the pre-medieval period. Such strong negative selection resulted in a quick loss of the mixed coat right after the domestication of modern horses (see our estimate for the underlying frequency trajectory of the mixed coat in Figure S5l). All are consistent with those shown in Figure 3.

## 4. Discussion

To overcome the key shortcoming of He et al. (2022), which does not aim to consider genetic interactions, in this work we have introduced a novel Bayesian procedure for inferring temporally variable selection from the data on aDNA sequences with the flexibility of modelling linkage and epistasis. We build our method upon the two-layer HMM framework of He et al. (2022), thereby enabling sample uncertainty arising from the damage and fragmentation of aDNA molecules. To allow genetic interactions, we introduce a Wright-Fisher diffusion to characterise the underlying evolutionary dynamics of two linked genes under phenotypic selection, which is modelled through the differential fitness of different phenotypic traits with a genotype-phenotype map. To ensure computational efficiency, we carry out the posterior computation with a robust adaptive PMMH algorithm, where unlike the original PMMH of Andrieu et al. (2010), we adopt the adaptation strategy of Vihola (2012) to tune the covariance structure of the proposal to achieve a coerced acceptance rate in our procedure. Furthermore, our approach permits the reconstruction of the underlying population gamete frequency trajectories and provides the flexibility of modelling time-varying demographic histories.

We have analysed horse pigmentation loci, e.g., *ASIP* and *MC1R* associated with base coat colours and *KIT13* and *KIT16* associated with pinto coat patterns, based on the ancient horse samples from previous studies of Ludwig et al. (2009), Pruvost et al. (2011) and Wutke et al. (2016). Our results illustrate that black, chestnut, tobiano and sabino horses were all favoured by selection from the domestication of modern horses, but selection acting on horse coat colours and patterns changed in the Middle Ages, which could be resulted from shifts in uses, cultures and religions. For example, our results provide strong evidence that tobiano horses experienced positive selection in the pre-medieval period but negative selection in the medieval period. This can be explained as resulting in part from that spotting could be the mark used to distinguish wild horses from domestic horses after modern horses were domesticated, but that differentiation might be no longer necessary as wild horses became scarcer and extinct. From historical records, we also find that initially spotted horses were associated with positive cultural image but had a negative connotation after a series of epidemics in the Middle Ages, which could have significant consequences on their popularity. In addition, the advent of weaponry in the Middle Ages could influence the preference of spotted horses since they were an easier target than solid horses. See Wutke et al. (2016) and references therein.

We have run additional simulations to test our procedure (see File S2 for simulation settings and results). Our results reveal that our approach can accurately estimate selection coefficients and discriminate whether selection acts on the phenotypic trait and whether selection changes at the given time (see Figures S6–S9 and Tables S9–S12, where we show the results produced with simulations that mimicked the ancient horse samples but in more general settings). Although we test our method with the demographic histories shown in Figure S1 and the genotype-phenotype maps described in Tables 1 and 3, in principle, the conclusions we have drawn here hold for other demographic histories and genotype-phenotype maps. Our simulation studies also indicate that misspecifying demographic histories can greatly alter the inference of selection (see Figures S10 and S11 and Tables S13 and S14), and therefore a method that can jointly estimate demographic and selective parameters is anticipated to boost the power, which can be an important topic of future research. Although our work infers selection on phenotypes, selection on genotypes can be measured if we assume 10 distinct phenotypes. Unlike He et al. (2020b), where they defined the fitness values of single-locus genotypes with a selection coefficient and a dominance parameter and assumed the fitness values of two-locus genotypes to be determined multiplicatively from those at individual loci, our method is built upon a general model of selection, therefore enabling *e.g*., heterozygote advantage selection.

To show the improvement attainable by modelling genetic interactions, we plot the resulting posteriors for *ASIP* and *MC1R* in Figure 4 and *KIT13* and *KIT16* in Figure 5, respectively, which are produced through the approach of He et al. (2022) with the same settings as adopted in our adaptive PMMH algorithm. We summarise the results for base coat colours and pinto coat patterns with their 95% HPD intervals in Tables S7 and S8, respectively. Moreover, additional simulation studies are left in File S2 to further show the improvement resulting from modelling linkage and epistasis (see Figures S12 and S13 and Tables S15 and S16).

**Figure 4:**
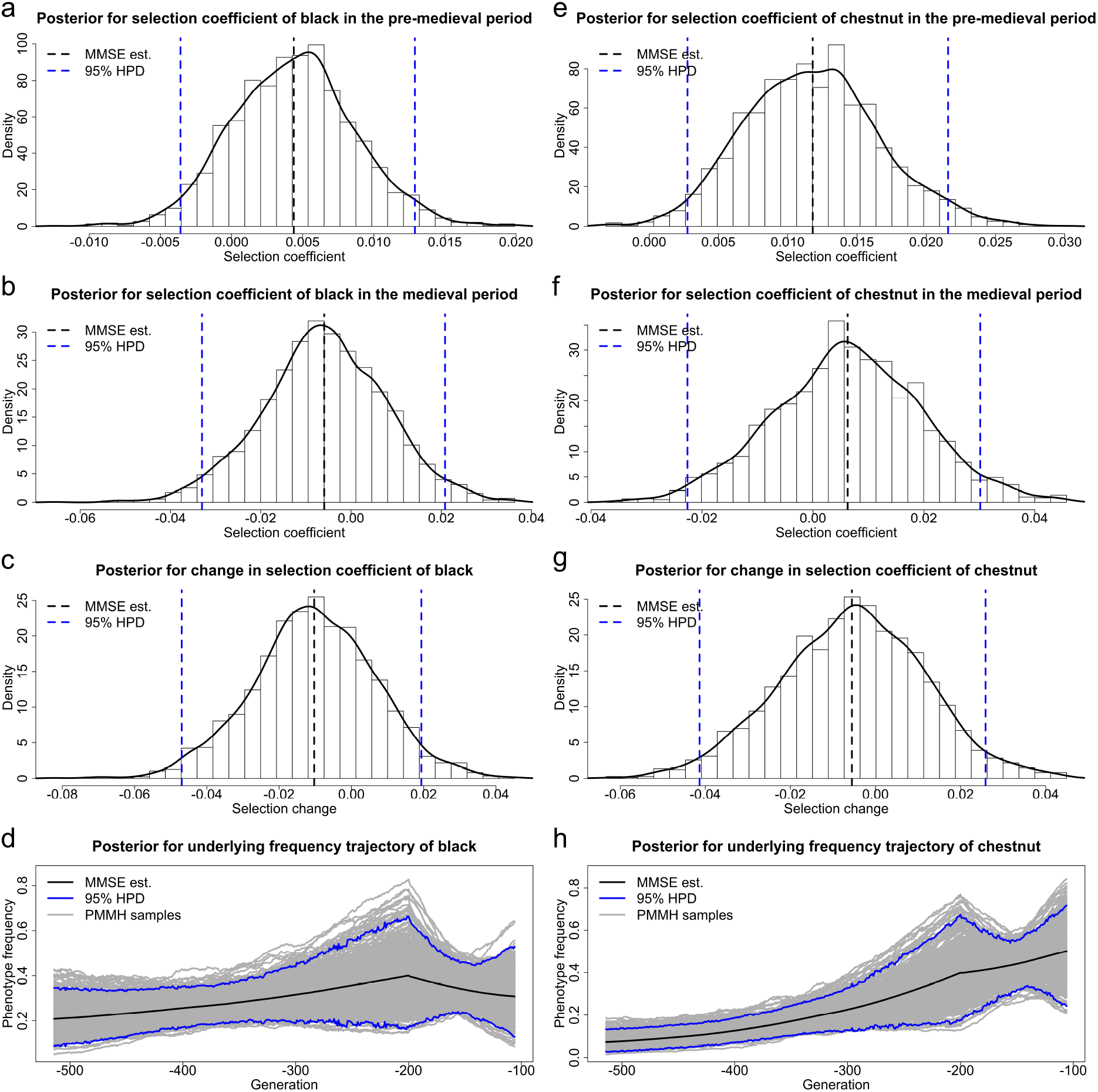
Posteriors for selection of base coat colours in the pre-medieval and medieval period and underlying frequency trajectories of each phenotypic trait in the population produced through the method of He et al. (2022), (a)-(d) for the black coat and (e)-(h) for the chestnut coat, respectively.

**Figure 5:**
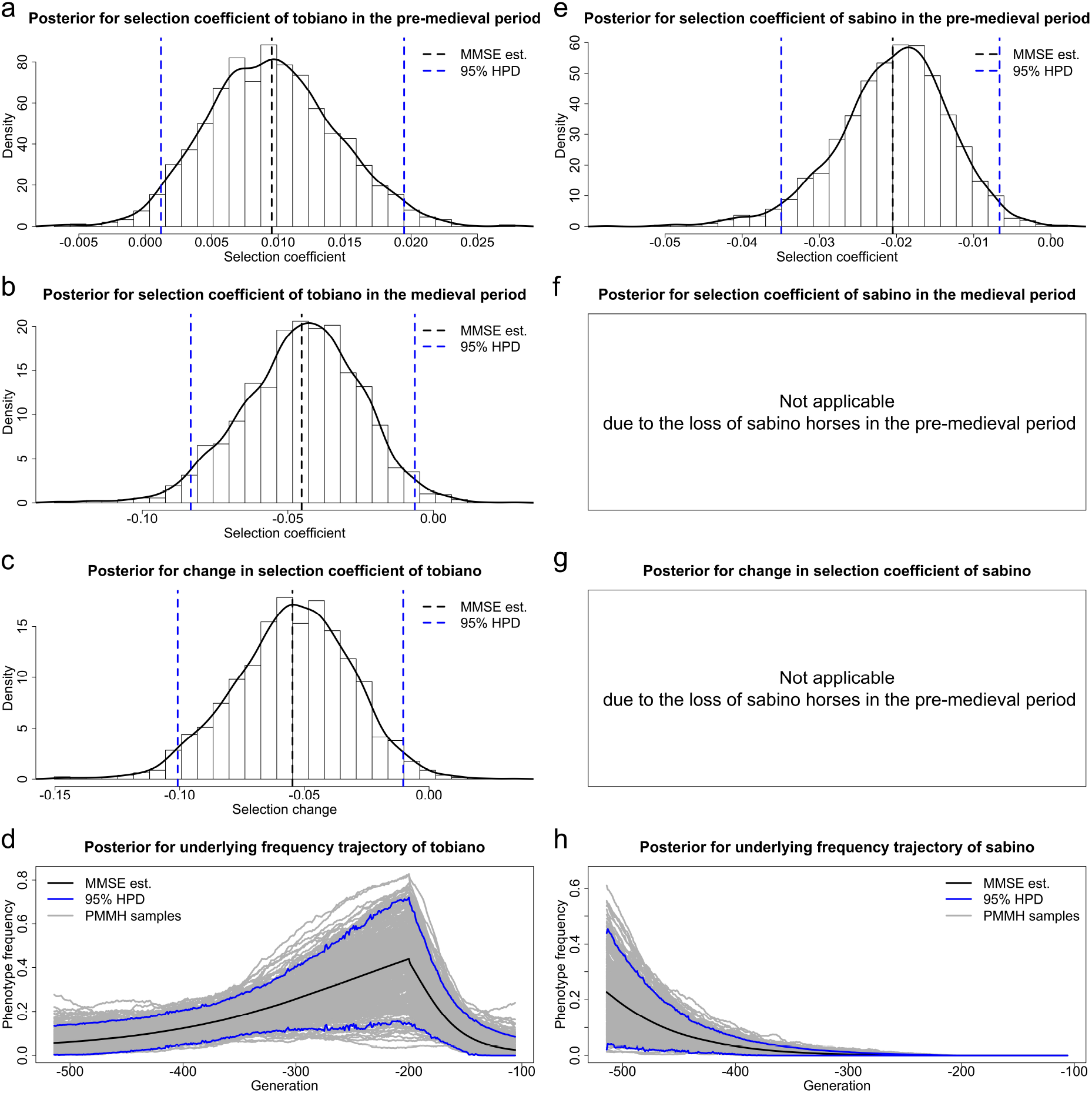
PPosteriors for selection of pinto coat patterns in the pre-medieval and medieval period and underlying frequency trajectories of each phenotypic trait in the population produced through the method of He et al. (2022), (a)-(d) for the tobiano coat and (e)-(h) for the sabino coat, respectively.

The results for *ASIP* and *MC1R* produced by the method of He et al. (2022) are consistent with those illustrated in Figure 2 except for selection acting on the black coat in the medieval period. Both results suggest that in the medieval period the black coat was negatively selected, but its intensity (i.e., 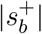) estimated by the approach of He et al. (2022) is substantially lower than that produced by our method. This is because ignoring epistasis causes the *aa/ee* genotype to be incorrectly attributed to the black coat, and most samples with *aa/ee* are medieval horses. The results for *KIT13* and *KIT16* produced by the approach of He et al. (2022) are compatible with those shown in Figure 3 except for selection acting on the sabino coat in the pre-medieval period. As shown in Figure 5, sabino horses experienced negative selection from modern horse domestication until extinction (in approximately AD 200), suggesting that a continuous decrease in sabino horses resulted from negative selection. However, we observe from Figure 3 that sabino horses were positively selected in the pre-medieval period but with a continuous decline in their frequency until extinction when linkage is modelled. This drop was probably triggered by sabino horses being somewhat out-competed by tobiano horses.

For aDNA data with high degrees of uncertainty, working on genotype likelihoods rather than called genotypes has been illustrated to be beneficial in the inference of selection (He et al., 2022). Genotype likelihoods are typically calculated with the aligned reads and associated mapping and sequencing quality scores, which facilitates the incorporation of uncertainty regarding genotypes arising from base calling, alignment and assembly (Nielsen et al., 2011). Recent methodological advances in genotype imputation, in particular those designed for imputing low-coverage data (*e.g*., Hui et al., 2020; Rubinacci et al., 2021; Ausmees & Nettelblad, 2023), present a possibility to improve aDNA studies (da Mota et al., 2022). Our procedure can directly work on imputed genotypes (and posterior genotype probabilities produced in most imputation approaches) and can be naturally extended to allow for phased haplotypes by combining with He et al. (2020b). Note that our approach can only leverage sample uncertainty captured by genotype likelihoods. To better model sample uncertainty associated with aDNA like that introduced by post-mortem C>T and G>A deamination, an important consideration is to integrate the model for genotype likelihood calculations based on aDNA sequences, introduced by Kousathanas et al. (2017) or Renaud et al. (2019), into our HMM framework. A further assumption of this work, in common with other studies, is of temporal continuity of populations. As shown by Silva et al. (2017) and Ortega-Del Vecchyo & Slatkin (2019), this may be a strong assumption, and inferences may be confounded by a history of gene flow or population replacement. These effects can be addressed in future, using more general spatial models.

Our extension inherits the desirable features of He et al. (2022) with its key limitation that all samples are assumed to be drawn after the respective mutations occurring at both loci. Since allele age is usually unavailable, we have to restrict our inference to a certain time window, e.g., from the time after which the mutant alleles at both loci have been observed in the sample or the time before which we assume that the mutant alleles at both loci have already existed in the population, which however could dramatically bias the inference of selection. Backward-in-time simulation of the Wright-Fisher diffusion (Griffiths, 2003; Coop & Griffiths, 2004) is anticipated to circumvent this problem. Moreover, how to extend our method to handle multiple interacting genes (Terhorst et al., 2015; Sohail et al., 2021, 2022) and estimate selection coefficients with their timing of changes (Shim et al., 2016; Mathieson, 2020; Mathieson & Terhorst, 2022) will also be the topics of our future investigation.

## Acknowledgements

We thank the anonymous reviewers and the editor for their helpful comments on the earlier version of this work. This work was carried out using the computational facilities of the Advanced Computing Research Centre, University of Bristol-http://www.bristol.ac.uk/acrc/.

## Data Accessibility Statement

The authors state that all data necessary for confirming the conclusions of the present work are represented completely within the article. Source code implementing the adaptive version of the PMMH algorithm described in this work is available at https://github.com/zhangyi-he/WFM-2L-DiffusApprox-AdaptPMMH/, where the standard version of the PMMH algorithm is also available.

## Author Contributions

Z.H. designed the project and developed the method; Z.H., X.D. and W.L. implemented the method; X.D. and W.L. analysed the data under the supervision of Z.H., M.B. and F.Y.; Z.H. wrote the manuscript; X.D., W.L., M.B. and F.Y. reviewed the manuscript.

## Notes

### Competing Interest Statement

The authors have declared no competing interest.

### Summary of Updates

We have reanalysed the aDNA data from horse pigmentation loci, where we excluded the ancient horse samples older than 4200 years ago (the domestication of modern horses). We performed the inference of selection that acts on the base coat colour controlled by ASIP and MC1R and the pinto coat pattern determined by KIT13 and KIT16 and tested the hypothesis that there was a shift in human preferences for horse coat colours and patterns in the Middle Ages when horses began to be differentiated by use.

