## Supporting Information for "Estimating temporally variable selection intensity from ancient DNA data with the flexibility of modelling linkage and epistasis"

### File S1. Additional results for the analysis of pigmentation loci in ancient horses

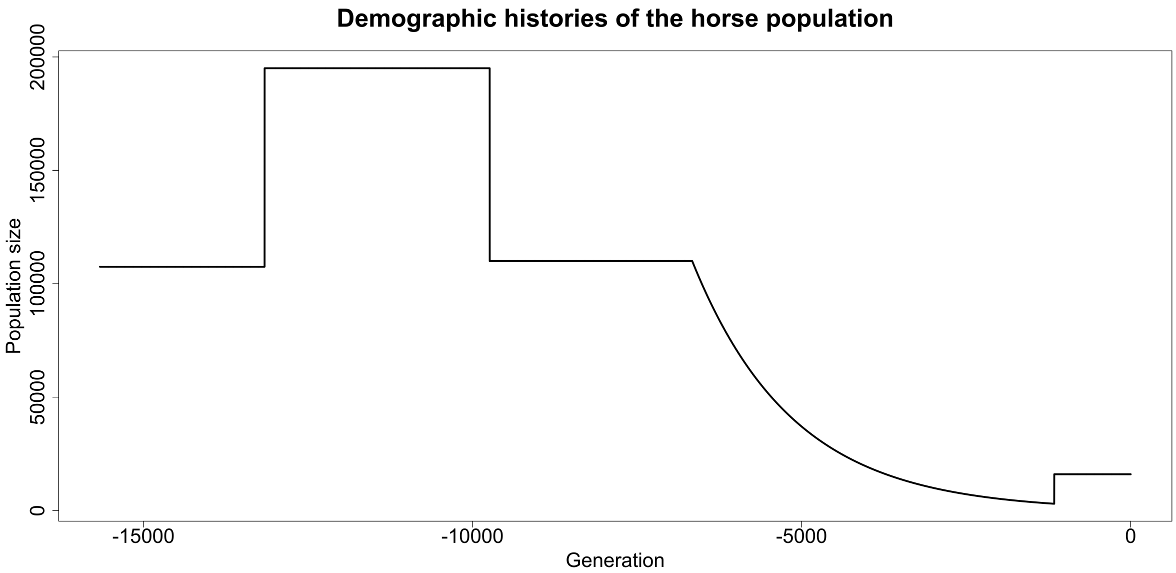


Figure S1: Demographic histories of the horse population reported in Der Sarkissian et al. (2015).

| Coat | Sel. coeff. | MMSE est. | 95% HPD | Prob. for -ve. | Prob. for +ve. |
|---|---|---|---|---|---|
| black | $s_b^-$ | 0.00551 | [−0.00333, 0.01509] | 0.1205 | 0.8795 |
| | $s_b^+$ | −0.00972 | [−0.04749, 0.03255] | 0.6660 | 0.3340 |
| | $\Delta s_b$ | −0.01522 | [−0.06508, 0.02925] | 0.7275 | 0.2725 |
| chestnut | $s_c^-$ | 0.01326 | [ 0.00280, 0.02269] | 0.0020 | 0.9980 |
| | $s_c^+$ | 0.00380 | [−0.02535, 0.03361] | 0.3870 | 0.6130 |
| | $\Delta s_c$ | −0.00946 | [−0.04470, 0.02941] | 0.6975 | 0.3025 |

Table S2: MMSE estimates of the selection coefficients and their changes for horse base coat colours with their 95% HPD intervals, as well as posterior probabilities for negative selection/change and positive selection/change, corresponding to Figure 2. Prob. for -ve. and +ve. stand for posterior probabilities for negative selection/change and positive selection/change, respectively.

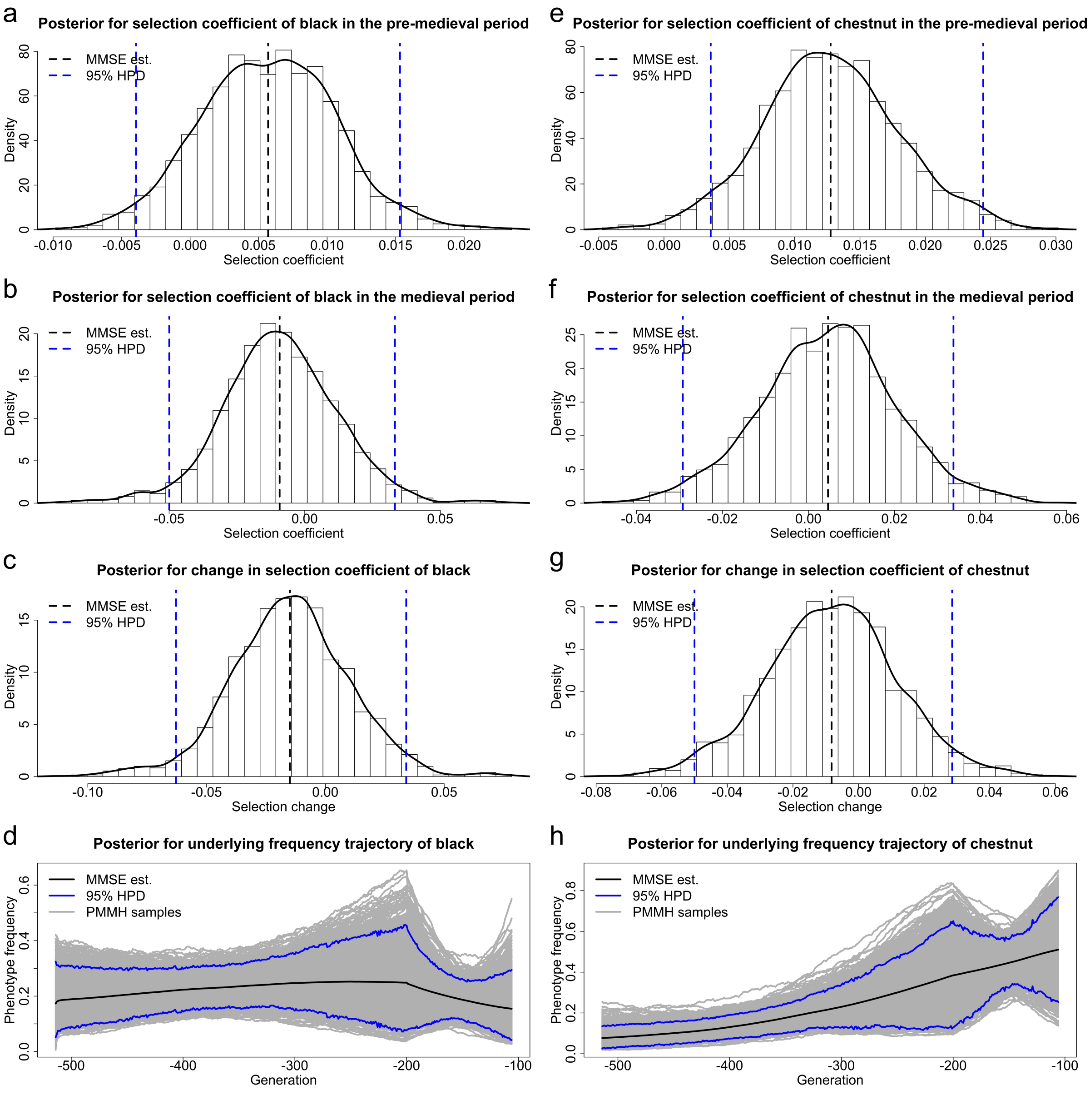


Figure S2: Posteriors for selection of base coat colours in the pre-medieval and medieval period and underlying frequency trajectories of each phenotypic trait in the population produced with the flat Dirichlet prior for the starting population gamete frequencies, (a)-(d) for the black coat and (e)-(h) for the chestnut coat, respectively.

| Coat | Sel. coeff. | MMSE est. | 95% HPD | | Prob. for -ve. | Prob. for +ve. |
|---|---|---|---|---|---|---|
| black | $s_b^-$ | 0.00569 | [−0.00395, | 0.01532] | 0.1285 | 0.8715 |
| | $s_b^+$ | −0.00924 | [−0.04997, | 0.03326] | 0.6825 | 0.3175 |
| | $\Delta s_b$ | −0.01494 | [−0.06287, | 0.03401] | 0.7430 | 0.2570 |
| chestnut | $s_c^-$ | 0.01275 | [ 0.00358, | 0.02443] | 0.0070 | 0.9930 |
| | $s_c^+$ | 0.00455 | [−0.02922, | 0.03371] | 0.3815 | 0.6185 |
| | $\Delta s_c$ | −0.00821 | [−0.05002, | 0.02853] | 0.6600 | 0.3400 |

Table S3: MMSE estimates of the selection coefficients and their changes for horse base coat colours with their 95% HPD intervals, as well as posterior probabilities for negative selection/change and positive selection/change, corresponding to Figure S2. Prob. for -ve. and +ve. stand for posterior probabilities for negative selection/change and positive selection/change, respectively.

| Coat | Sel. coeff. | MMSE est. | 95% HPD | Prob. for -ve. | Prob. for +ve. |
|---|---|---|---|---|---|
| tobiano | $s_{to}^{-}$ | 0.01967 | $[\ 0.00835, \ 0.03188]$ | 0.0000 | 1.0000 |
| | $s_{to}^{+}$ | $-0.06462$ | $[-0.10543, -0.02405]$ | 1.0000 | 0.0000 |
| | $\Delta s_{to}$ | $-0.08429$ | $[-0.13449, -0.03643]$ | 1.0000 | 0.0000 |
| sabino | $s_{sb}^{-}$ | 0.03214 | $[-0.00610, \ 0.08143]$ | 0.0595 | 0.9405 |
| | $s_{sb}^{+}$ | N/A | N/A | N/A | N/A |
| | $\Delta s_{sb}$ | N/A | N/A | N/A | N/A |

Table S4: MMSE estimates of the selection coefficients and their changes for horse pinto coat patterns with their 95% HPD intervals, as well as posterior probabilities for negative selection/change and positive selection/change, corresponding to Figure 3. Prob. for -ve. and +ve. stand for posterior probabilities for negative selection/change and positive selection/change, respectively.

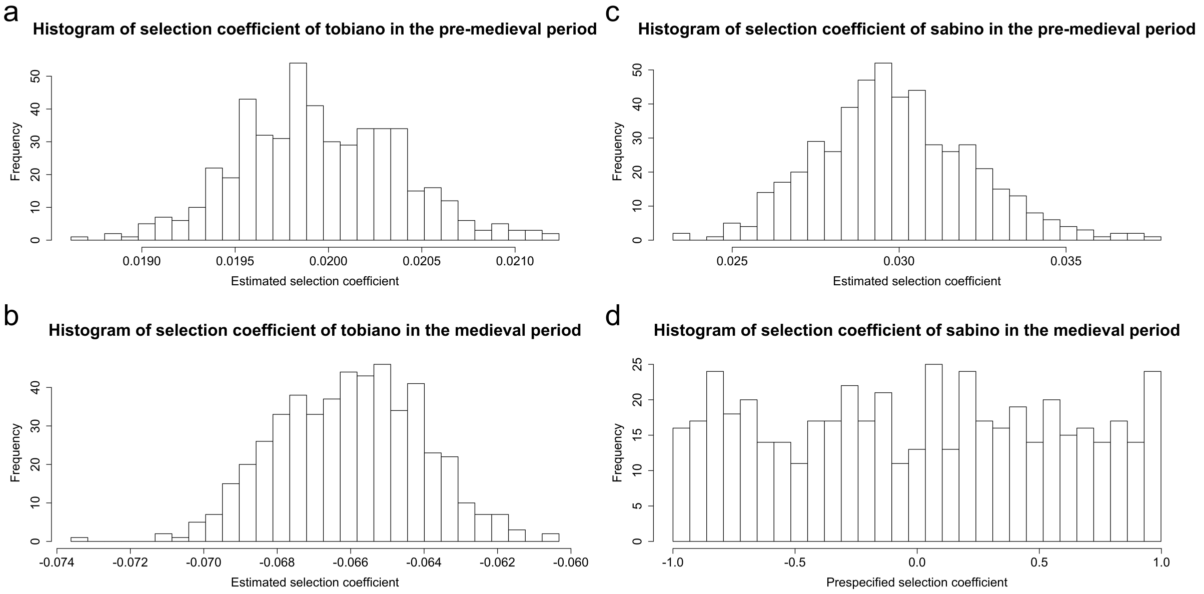


Figure S3: Empirical distributions of the estimates of the selection coefficients $s_{to}^{-}$, $s_{to}^{+}$ and $s_{sb}^{-}$ over 500 replicates that we run our procedure on the ancient horse samples with different prespecified values of the selection coefficient $s_{sb}^{+}$ uniformly drawn from $[-1, 1]$. The mean estimates of the selection coefficients $s_{to}^{-}$, $s_{to}^{+}$ and $s_{sb}^{-}$ are 0.02000, $-0.06602$ and 0.02990 with their standard deviation 0.00043, 0.00197 and 0.00234.

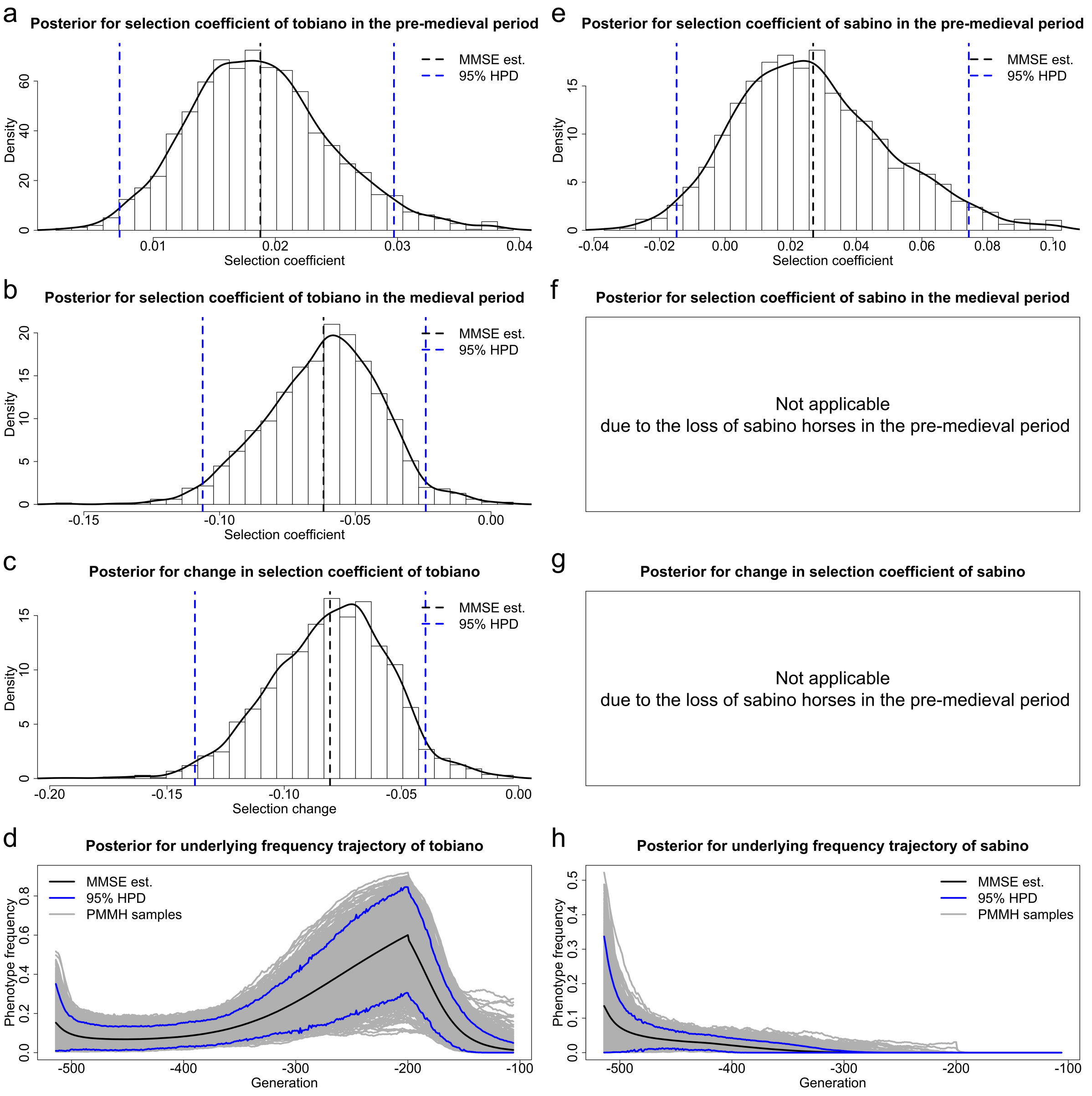


Figure S4: Posteriors for selection of pinto coat patterns in the pre-medieval and medieval period and underlying frequency trajectories of each phenotypic trait in the population produced with the flat Dirichlet prior for the starting population gamete frequencies, (a)-(d) for the tobiano coat and (e)-(h) for the sabino coat, respectively.

| Coat | Sel. coeff. | MMSE est. | 95% HPD | Prob. for -ve. | Prob. for +ve. |
|---|---|---|---|---|---|
| tobiano | $s_{to}^{-}$ | 0.01877 | [ 0.00725, 0.02971] | 0.0000 | 1.0000 |
| | $s_{to}^{+}$ | −0.06170 | [−0.10623, −0.02404] | 0.9980 | 0.0020 |
| | $\Delta s_{to}$ | −0.08047 | [−0.13808, −0.03976] | 1.0000 | 0.0000 |
| sabino | $s_{sb}^{-}$ | 0.02685 | [−0.01478, 0.07431] | 0.1085 | 0.8915 |
| | $s_{sb}^{+}$ | N/A | N/A | N/A | N/A |
| | $\Delta s_{sb}$ | N/A | N/A | N/A | N/A |

Table S5: MMSE estimates of the selection coefficients and their changes for horse pinto coat patterns with their 95% HPD intervals, as well as posterior probabilities for negative selection/change and positive selection/change, corresponding to Figure S4. Prob. for -ve. and +ve. stand for posterior probabilities for negative selection/change and positive selection/change, respectively.

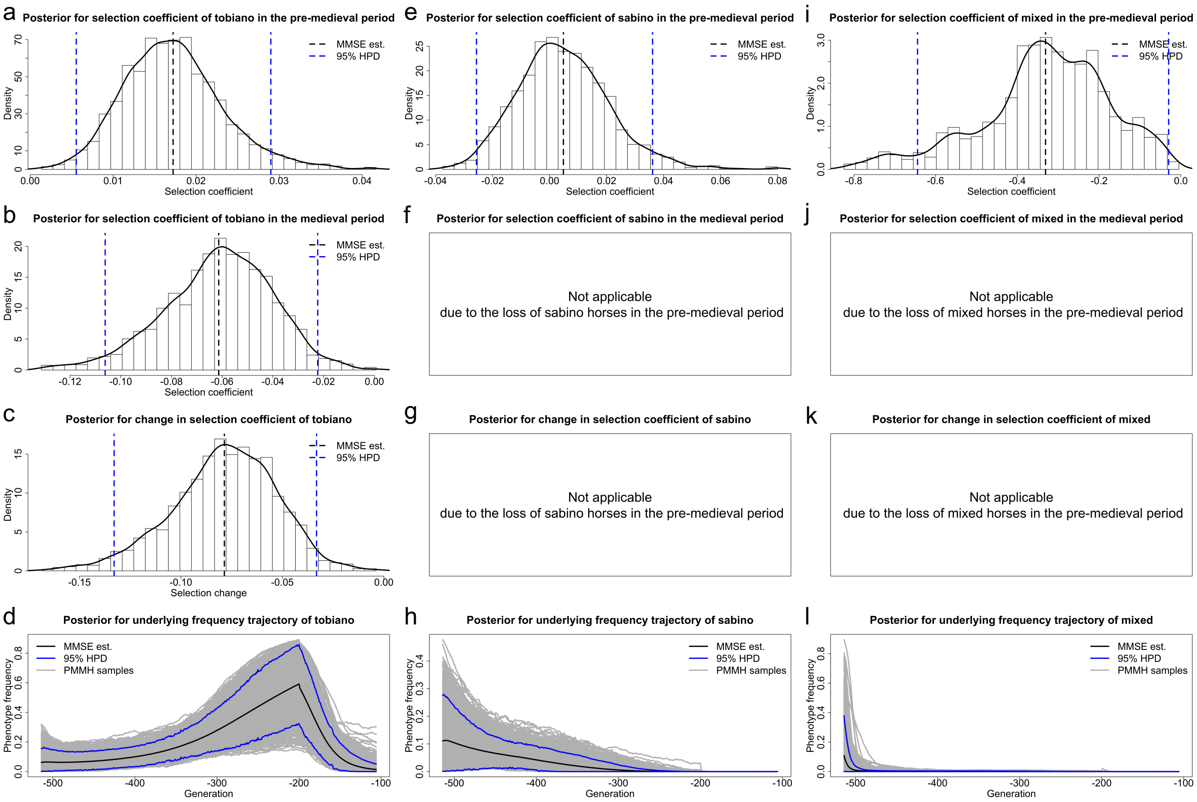


Figure S5: Posteriors for selection of pinto coat patterns in the pre-medieval and medieval period and underlying frequency trajectories of each phenotypic trait in the population, (a)-(d) for the tobiano coat, (e)-(h) for the sabino coat and (i)-(l) for the mixed coat, respectively.

| Coat | Sel. coeff. | MMSE est. | 95% HPD | Prob. for -ve. | Prob. for +ve. |
|---|---|---|---|---|---|
| tobiano | $s_{to}^-$ | 0.01722 | [ 0.00555, 0.02900] | 0.0000 | 1.0000 |
| | $s_{to}^+$ | −0.06138 | [−0.10615, −0.02235] | 0.9985 | 0.0015 |
| | $\Delta s_{to}$ | −0.07860 | [−0.13294, −0.03318] | 1.0000 | 0.0000 |
| sabino | $s_{sb}^-$ | 0.00483 | [−0.02565, 0.03613] | 0.4045 | 0.5955 |
| | $s_{sb}^+$ | N/A | N/A | N/A | N/A |
| | $\Delta s_{sb}$ | N/A | N/A | N/A | N/A |
| mixed | $s_{mx}^-$ | −0.33106 | [−0.64419, −0.02992] | 1.0000 | 0.0000 |
| | $s_{mx}^+$ | N/A | N/A | N/A | N/A |
| | $\Delta s_{mx}$ | N/A | N/A | N/A | N/A |

Table S6: MMSE estimates of the selection coefficients and their changes for horse pinto coat patterns with their 95% HPD intervals, as well as posterior probabilities for negative selection/change and positive selection/change, corresponding to Figure S5. Prob. for -ve. and +ve. stand for posterior probabilities for negative selection/change and positive selection/change, respectively.

| Coat. | Sel. coeff. | MMSE est. | 95% HPD | Prob. for -ve. | Prob. for +ve. |
|---|---|---|---|---|---|
| black | $s_b^-$ | 0.00437 | $[-0.00359, \quad 0.01289]$ | 0.1590 | 0.8410 |
| | $s_b^+$ | $-0.00594$ | $[-0.03302, \quad 0.02084]$ | 0.6805 | 0.3195 |
| | $\Delta s_b$ | $-0.01031$ | $[-0.04696, \quad 0.01937]$ | 0.7305 | 0.2695 |
| chestnut | $s_c^-$ | 0.01181 | $[\ \ 0.00276, \quad 0.02158]$ | 0.0035 | 0.9965 |
| | $s_c^+$ | 0.00626 | $[-0.02262, \quad 0.03017]$ | 0.3065 | 0.6935 |
| | $\Delta s_c$ | $-0.00554$ | $[-0.04135, \quad 0.02588]$ | 0.6265 | 0.3735 |

Table S7: MMSE estimates of the selection coefficients and their changes for horse base coat colours with their 95% HPD intervals, as well as posterior probabilities for negative selection/change and positive selection/change, corresponding to Figure 4. Prob. for -ve. and +ve. stand for posterior probabilities for negative selection/change and positive selection/change, respectively.

| Coat | Sel. coeff. | MMSE est. | 95% HPD | Prob. for -ve. | Prob. for +ve. |
|---|---|---|---|---|---|
| tobiano | $s_{to}^{-}$ | 0.00952 | [ 0.00118, 0.01948] | 0.0155 | 0.9845 |
| | $s_{to}^{+}$ | −0.04526 | [−0.08336, −0.00637] | 0.9915 | 0.0085 |
| | $\Delta s_{to}$ | −0.05478 | [−0.10080, −0.01037] | 0.9925 | 0.0075 |
| sabino | $s_{sb}^{-}$ | −0.02049 | [−0.03494, −0.00664] | 0.9990 | 0.0010 |
| | $s_{sb}^{+}$ | N/A | N/A | N/A | N/A |
| | $\Delta s_{sb}$ | N/A | N/A | N/A | N/A |

Table S8: MMSE estimates of the selection coefficients and their changes for horse pinto coat patterns with their 95% HPD intervals, as well as posterior probabilities for negative selection/change and positive selection/change, corresponding to Figure 5. Prob. for -ve. and +ve. stand for posterior probabilities for negative selection/change and positive selection/change, respectively.

**File S2. Simulation studies on performance evaluation**

We run extensive simulations to achieve empirical results on the performance of our method described in this work in two scenarios, horse base coat colours (*i.e.*, two genes with epistasis) and pinto coat patterns (*i.e.*, two genes with linkage), respectively. More specifically, we generate the underlying frequency trajectories of the gametes in the population through forward-in-time simulations of the Wright-Fisher model in Eq. (1). Each individual in the sample is then drawn from the population by multinomial sampling in Eq. (7). Given the genotype of each sampled individual, the corresponding genotype likelihoods at each locus are independently produced by Dirichlet sampling like He et al. (2022). For clarity, we write down the procedure that we follow to generate datasets of genotype likelihoods:

Step 1: Generate $s_1^-$, $s_1^+$, $s_2^-$, $s_2^+$ and $\boldsymbol{x}_{1:K}$.

- Step 1a: Draw $s_i^- \sim \text{Uniform}(-0.05, 0.05)$ with a probability of 0.5 otherwise set $s_i^- = 0$ for $i = 1, 2$.
- Step 1b: Draw $s_i^+ \sim \text{Uniform}(-0.05, 0.05)$ with a probability of 0.5 otherwise set $s_i^+ = s_i^-$ for $i = 1, 2$.
- Step 1c: Draw $\boldsymbol{x}_1$ by following the procedure described in Section 3.
- Step 1d: Draw $\boldsymbol{x}_{1:K}$ by simulating the Wright-Fisher model in Eq. (1) with $s_1^-$, $s_1^+$, $s_2^-$, $s_2^+$ and $\boldsymbol{x}_1$.

Repeat Step 2 for $k = 1, 2, \ldots, K$:

Step 2: Generate $p(\boldsymbol{r}_{l,n,k} \mid g)$ for $g = 0, 1, 2$, $l = 1, 2$ and $n = 1, 2, \ldots, N_k$:

- Step 2a: Draw $\boldsymbol{g}_{n,k} \sim p(\boldsymbol{g} \mid \boldsymbol{x}_k)$ in Eq. (7).
- Step 2b: Draw $(p(\boldsymbol{r}_{l,n,k} \mid g = 0), p(\boldsymbol{r}_{l,n,k} \mid g = 1), p(\boldsymbol{r}_{l,n,k} \mid g = 2)) \sim \text{Dirichlet}(\boldsymbol{\alpha}^{l,n,k})$ for $l = 1, 2$, where $\boldsymbol{\alpha}^{l,n,k} = (\alpha_0^{l,n,k}, \alpha_1^{l,n,k}, \alpha_2^{l,n,k})$.

We take the population size to be the same as those adopted in Section 3. As in He et al. (2022), we take the parameter $\alpha_{g_{l,n,k}}^{l,n,k}$ to be $\phi\psi$ and the other two to be $(1 - \phi)\psi/2$ with $\phi = 0.95$ and $\psi = 1$, which gives rise to an average missing rate of 11.1% and an average error rate of 0.5% with a common threshold for genotype calling (*i.e.*, 10 times more likely, see Kim et al., 2011). We adopt the same sampling scheme (*i.e.*, sample sizes and times) and event time as the aDNA dataset of horses (see Table S1).

For each scenario, we repeatedly run the procedure described above until we obtain 500 simulated datasets, in each of which we can observe at least one individual with a mutant phenotype in the sample drawn in a pre- and post-event period, respectively. We run our adaptive PMMH procedure on each simulated dataset with the same settings as those adopted in Section 3 for the aDNA dataset of horses. We plot the empirical distributions of the resulting estimates of the selection coefficients for horse base coat colours and pinto coat patterns, respectively, with the receiver operating characteristic (ROC) curves for identifying selection signatures and testing selection changes. The ROC curve is produced by plotting the true-positive rate (TPR) against the false-positive rate (FPR), where *e.g.*, the TPR and FPR are computed for each value of the posterior probability for being selected that is set to be a threshold to classify a phenotypic trait as experiencing selection. We also calculate the area under the ROC curve (AUC) to summarise the performance.

*Performance evaluation.* To test our method in the same settings as our analysis of pigmentation loci in ancient horses presented in Section 3, we generate 500 datasets of genotype likelihoods through the procedure described above, where we take the recombination rate to be the same as those adopted in each scenario in Section 3. For base coat colours, we take the coefficient of linkage disequilibrium to be $D = 0$ in Step 1c, and for pinto coat patterns, we take the selection coefficients of the mixed coat to be $s_{mx}^- = s_{mx}^+ = -1$ in Steps 1a and 1b. We illustrate the empirical distributions of the resulting estimates in Figure S6 for base coat colours and in Figure S7 for pinto coat patterns, respectively, along with the ROC curves and AUC values for detecting selection signatures and testing selection changes. See Tables S9 and S10 for the mean bias and root mean square error (RMSE) of the resulting estimates.

| Coat | Sel. coeff. | Bias | RMSE |
|---|---|---|---|
| black | $s_b^-$ | −0.00850 | 0.03170 |
| | $s_b^+$ | 0.00583 | 0.09876 |
| chestnut | $s_c^-$ | −0.00413 | 0.01732 |
| | $s_c^+$ | −0.00793 | 0.07327 |

Table S9: Mean bias and RMSE in MMSE estimates of the selection coefficients for horse base coat colours, corresponding to Figure S6. Mean bias and RMSE are calculated with 500 replicates.

To test our method in more general settings, we generate 500 datasets of genotype likelihoods through the procedure described above, where for each case we assume that the two loci reside on

| Coat | Sel. coeff. | Bias | RMSE |
|---|---|---|---|
| tobiano | $s_{to}^-$ | −0.01171 | 0.07460 |
| | $s_{to}^+$ | 0.00125 | 0.13792 |
| sabino | $s_{sb}^-$ | −0.01048 | 0.07612 |
| | $s_{sb}^+$ | −0.00520 | 0.12512 |

Table S10: Mean bias and RMSE in MMSE estimates of the selection coefficients for horse pinto coat patterns, corresponding to Figure S7. Mean bias and RMSE are calculated with 500 replicates.

the same chromosome, 50000 base pairs (bp) apart, with an average rate of recombination $10^{-8}$ crossover/bp. Notice that for pinto coat patterns, we uniformly draw the selection coefficients of the mixed coat $s_{mx}^-$ and $s_{mx}^+$ in Steps 1a and 1b like other selection coefficients and jointly estimate all selection coefficients. We show the empirical distributions of the resulting estimates in Figure S8 for base coat colours and in Figure S9 for pinto coat patterns, respectively, along with the ROC curves and AUC values for identifying selection signatures and testing selection changes. See Tables S11 and S12 for the mean bias and RMSE of the resulting estimates.

| Coat | Sel. coeff. | Bias | RMSE |
|---|---|---|---|
| black | $s_b^-$ | 0.00073 | 0.03594 |
| | $s_b^+$ | 0.00334 | 0.07558 |
| chestnut | $s_c^-$ | −0.00216 | 0.02669 |
| | $s_c^+$ | −0.00071 | 0.07300 |

Table S11: Mean bias and RMSE in MMSE estimates of the selection coefficients for horse base coat colours, corresponding to Figure S8. Mean bias and RMSE are calculated with 500 replicates.

| Coat | Sel. coeff. | Bias | RMSE |
|---|---|---|---|
| tobiano | $s_{to}^-$ | 0.02648 | 0.09650 |
| | $s_{to}^+$ | 0.01924 | 0.16859 |
| sabino | $s_{sb}^-$ | 0.02496 | 0.09051 |
| | $s_{sb}^+$ | 0.00432 | 0.18036 |
| mixed | $s_{mx}^-$ | 0.01907 | 0.09226 |
| | $s_{mx}^+$ | −0.00996 | 0.19032 |

Table S12: Mean bias and RMSE in MMSE estimates of the selection coefficients for horse pinto coat patterns, corresponding to Figure S9. Mean bias and RMSE are calculated with 500 replicates.

*Effect of misspecifying demographic histories.* To illustrate the influence caused by misspecifying demographic histories, we run our method on the datasets generated for Figures S6 and S7 with misspecified demographic histories (*i.e.*, we adopt the smallest horse population size of $N = 3000$, which is fixed over time). We show the empirical distributions of the resulting estimates in Figure S10 for base coat colours and in Figure S11 for pinto coat patterns, respectively, along with the ROC curves and AUC values for identifying selection signatures and testing selection changes. See Tables S13 and S14 for the mean bias and RMSE of the resulting estimates.

| Coat | Sel. coeff. | Bias | RMSE |
|---|---|---|---|
| black | $s_b^-$ | −0.01167 | 0.03629 |
| | $s_b^+$ | 0.00772 | 0.10789 |
| chestnut | $s_c^-$ | −0.00553 | 0.01857 |
| | $s_c^+$ | −0.00708 | 0.07079 |

Table S13: Mean bias and RMSE in MMSE estimates of the selection coefficients for horse base coat colours, corresponding to Figure S10. Mean bias and RMSE are calculated with 500 replicates for each gene.

| Coat | Sel. coeff. | Bias | RMSE |
|---|---|---|---|
| tobiano | $s_{to}^-$ | −0.01049 | 0.08708 |
| | $s_{to}^+$ | 0.02424 | 0.27115 |
| sabino | $s_{sb}^-$ | −0.00856 | 0.08043 |
| | $s_{sb}^+$ | 0.08658 | 0.64196 |

Table S14: Mean bias and RMSE in MMSE estimates of the selection coefficients for horse pinto coat patterns, corresponding to Figure S11. Mean bias and RMSE are calculated with 500 replicates for each gene.

*Effect of ignoring genetic interactions.* To illustrate the influence resulting from ignoring linkage and epistasis, we run the approach of He et al. (2022) on the datasets generated for Figures S6 and S7 (separately for each gene) with the same settings as those adopted in Section 4. We show the empirical distributions of the resulting estimates in Figure S12 for base coat colours and in Figure S13 for pinto coat patterns, respectively, along with the ROC curves and AUC values for identifying selection signatures and testing selection changes. See Tables S15 and S16 for the mean bias and RMSE of the resulting estimates.

| Coat | Sel. coeff. | Bias | RMSE |
|---|---|---|---|
| black | $s_b^-$ | −0.00788 | 0.03304 |
| | $s_b^+$ | 0.09694 | 2.04118 |
| chestnut | $s_c^-$ | −0.00392 | 0.01732 |
| | $s_c^+$ | 0.04530 | 1.21098 |

Table S15: Mean bias and RMSE in MMSE estimates of the selection coefficients for horse base coat colours, corresponding to Figure S12. Mean bias and RMSE are calculated with 500 replicates for each gene.

| Coat | Sel. coeff. | Bias | RMSE |
|---|---|---|---|
| tobiano | $s_{to}^{-}$ | − 0.03565 | 0.09758 |
| | $s_{to}^{+}$ | 8.91895 | 38.25395 |
| sabino | $s_{sb}^{-}$ | − 0.04356 | 0.12009 |
| | $s_{sb}^{+}$ | 11.20470 | 46.86521 |

Table S16: Mean bias and RMSE in MMSE estimates of the selection coefficients for horse pinto coat patterns, corresponding to Figure S13. Mean bias and RMSE are calculated with 500 replicates for each gene.

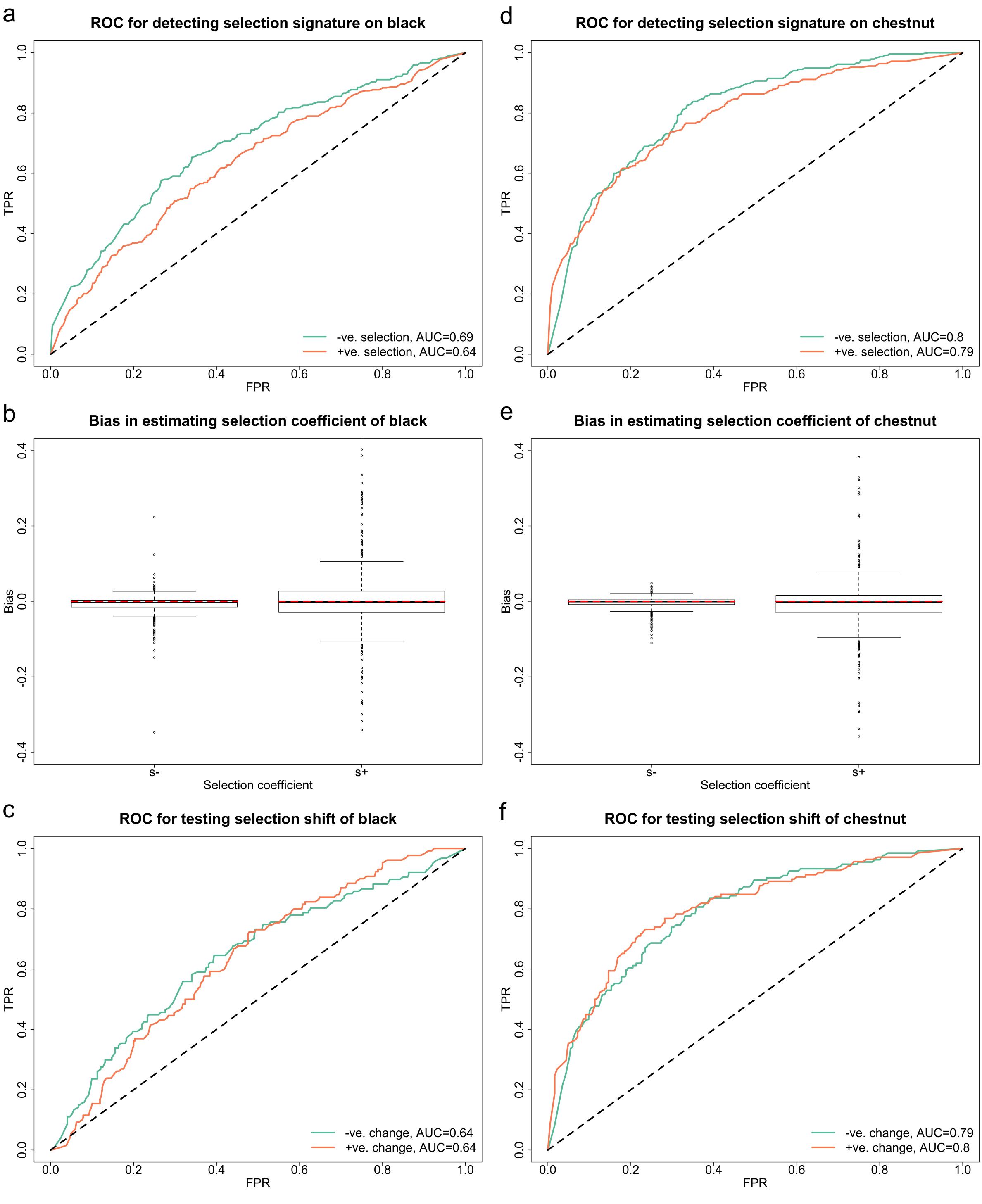


Figure S6: Empirical distributions for the bias in MMSE estimates of the selection coefficients for horse base coat colours with ROC curves and AUC values for detecting selection signatures and testing selection changes, (a)-(c) for the black coat and (d)-(f) for the chestnut coat, respectively. To aid visualisation, we remove the replicates in which the absolute value of bias is larger than 0.4 from (b) and (e), *i.e.*, 6 replicates from (b) and 0 replicates from (e), respectively.

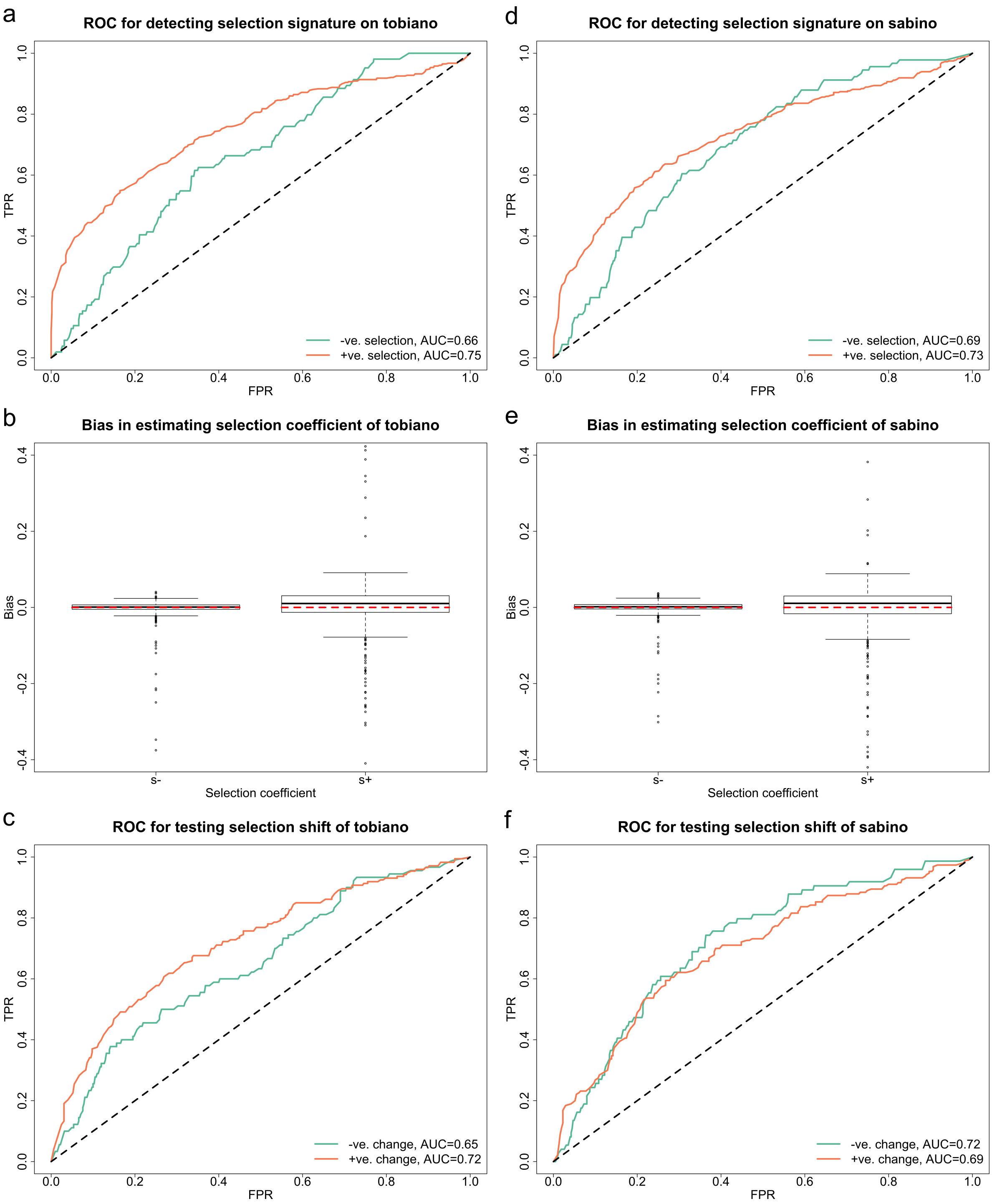


Figure S7: Empirical distributions for the bias in MMSE estimates of the selection coefficients for horse pinto coat patterns with ROC curves and AUC values for detecting selection signatures and testing selection changes, (a)-(c) for the tobiano coat and (d)-(f) for the sabino coat, respectively. To aid visualisation, we remove the replicates in which the absolute value of bias is larger than 0.4 from (b) and (e), *i.e.*, 31 replicates from (b) and 24 replicates from (e), respectively.

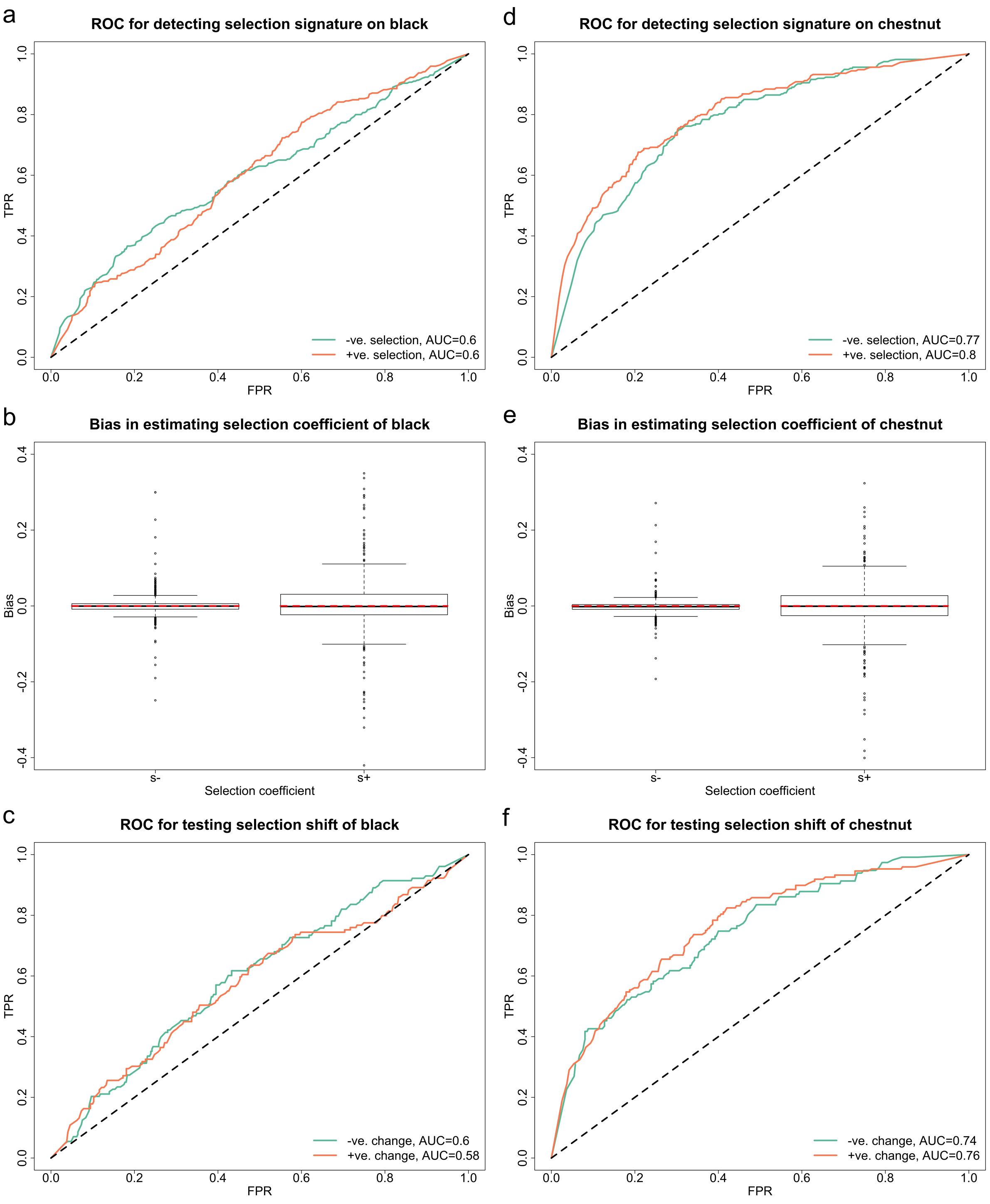


Figure S8: Empirical distributions for the bias in MMSE estimates of the selection coefficients for horse base coat colours with ROC curves and AUC values for detecting selection signatures and testing selection changes, (a)-(c) for the black coat and (d)-(f) for the chestnut coat, respectively. To aid visualisation, we remove the replicates in which the absolute value of bias is larger than 0.4 from (b) and (e), *i.e.*, 1 replicates from (b) and 2 replicates from (e), respectively.

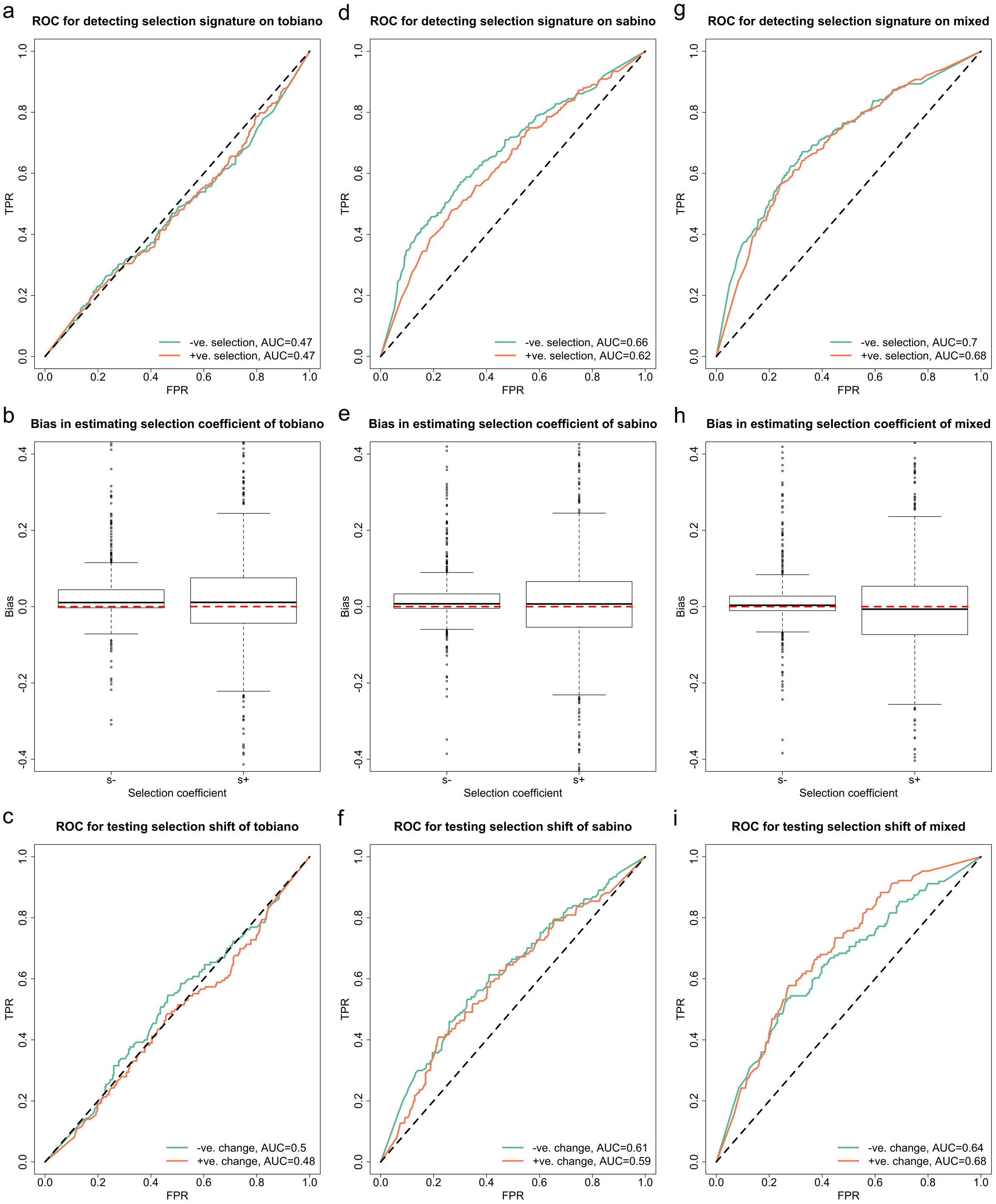


Figure S9: Empirical distributions for the bias in MMSE estimates of the selection coefficients for horse pinto coat patterns with ROC curves and AUC values for detecting selection signatures and testing selection changes, (a)-(c) for the tobiano coat, (d)-(f) for the sabino coat and (g)-(i) for the mixed coat, respectively. To aid visualisation, we remove the replicates in which the absolute value of bias is larger than 0.4 from (b), (e) and (h), *i.e.*, 33 replicates from (b), 34 replicates from (e) and 40 replicates from (h), respectively.

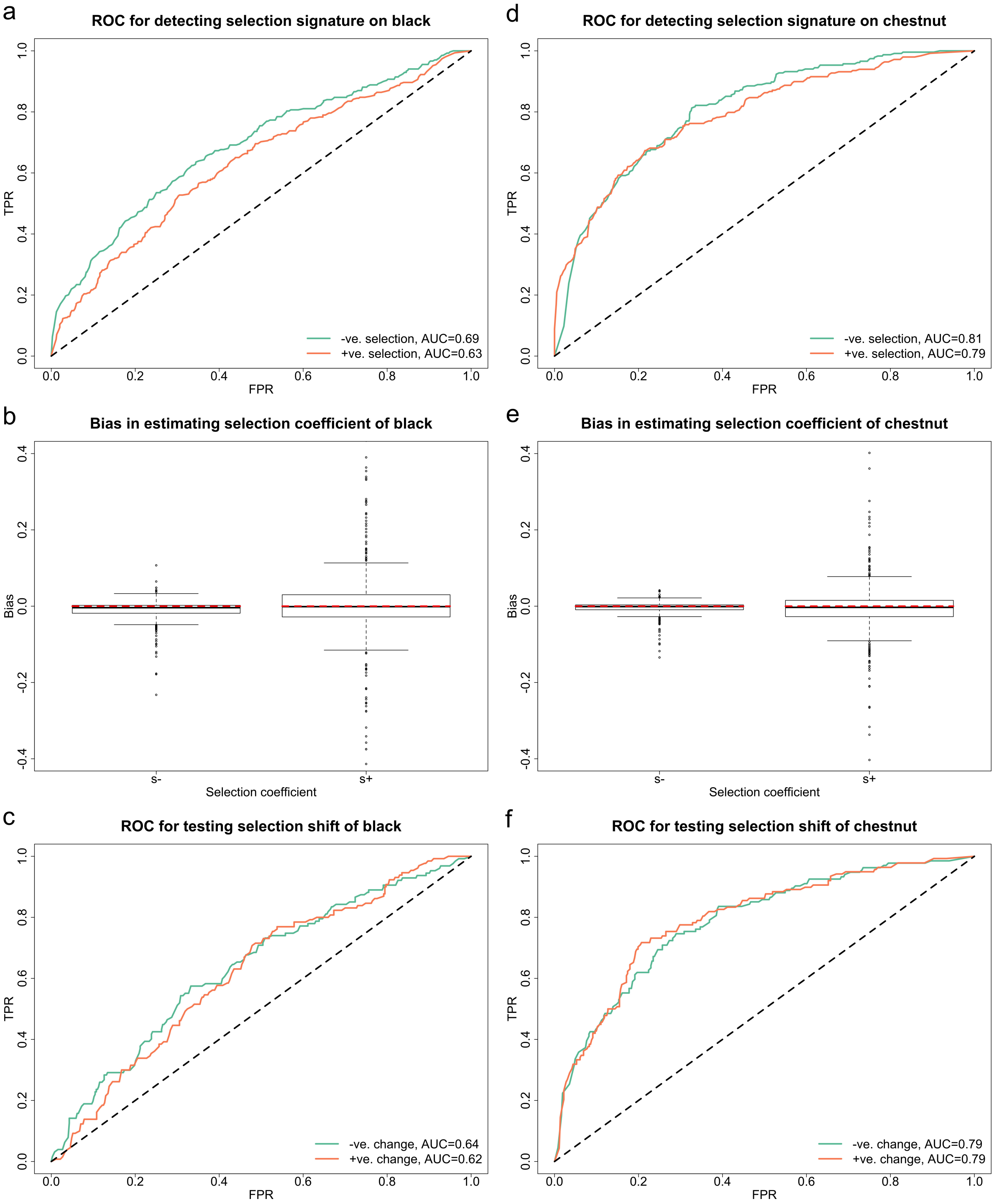


Figure S10: Empirical distributions for the bias in MMSE estimates of the selection coefficients for horse base coat colours with ROC curves and AUC values for detecting selection signatures and testing selection changes, (a)-(c) for the black coat and (d)-(f) for the chestnut coat, respectively. To aid visualisation, we remove the replicates in which the absolute value of bias is larger than 0.4 from (b) and (e), *i.e.*, 7 replicates from (b) and 2 replicates from (e), respectively.

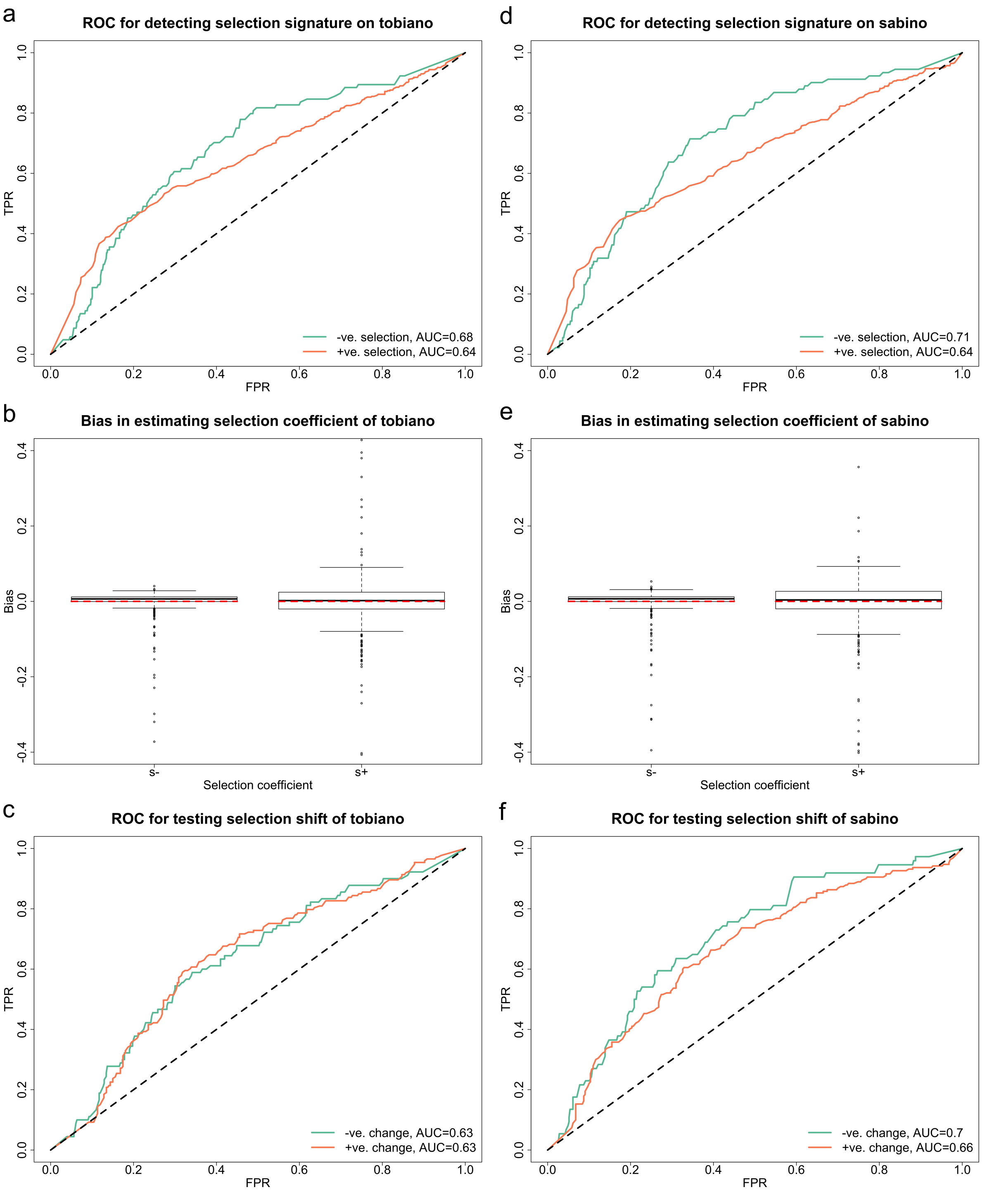


Figure S11: Empirical distributions for the bias in MMSE estimates of the selection coefficients for horse pinto coat patterns with ROC curves and AUC values for detecting selection signatures and testing selection changes, (a)-(c) for the tobiano coat and (d)-(f) for the sabino coat, respectively. To aid visualisation, we remove the replicates in which the absolute value of bias is larger than 0.4 from (b) and (e), *i.e.*, 40 replicates from (b) and 44 replicates from (e), respectively.

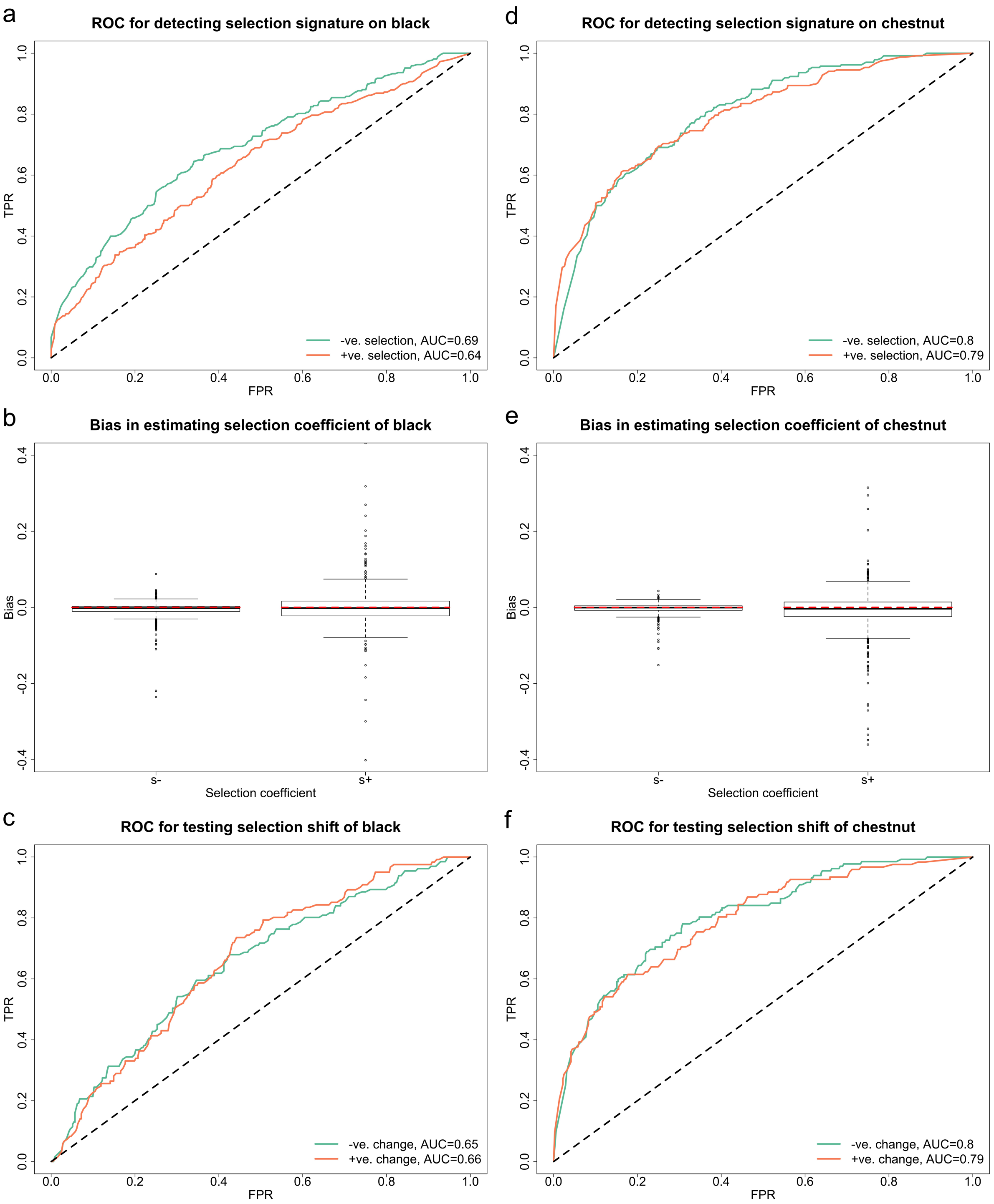


Figure S12: Empirical distributions for the bias in MMSE estimates of the selection coefficients for horse base coat colours with ROC curves and AUC values for detecting selection signatures and testing selection changes, (a)-(c) for the black coat and (d)-(f) for the chestnut coat, respectively. To aid visualisation, we remove the replicates in which the absolute value of bias is larger than 0.4 from (b) and (e), *i.e.*, 7 replicates from (b) and 1 replicates from (e), respectively.

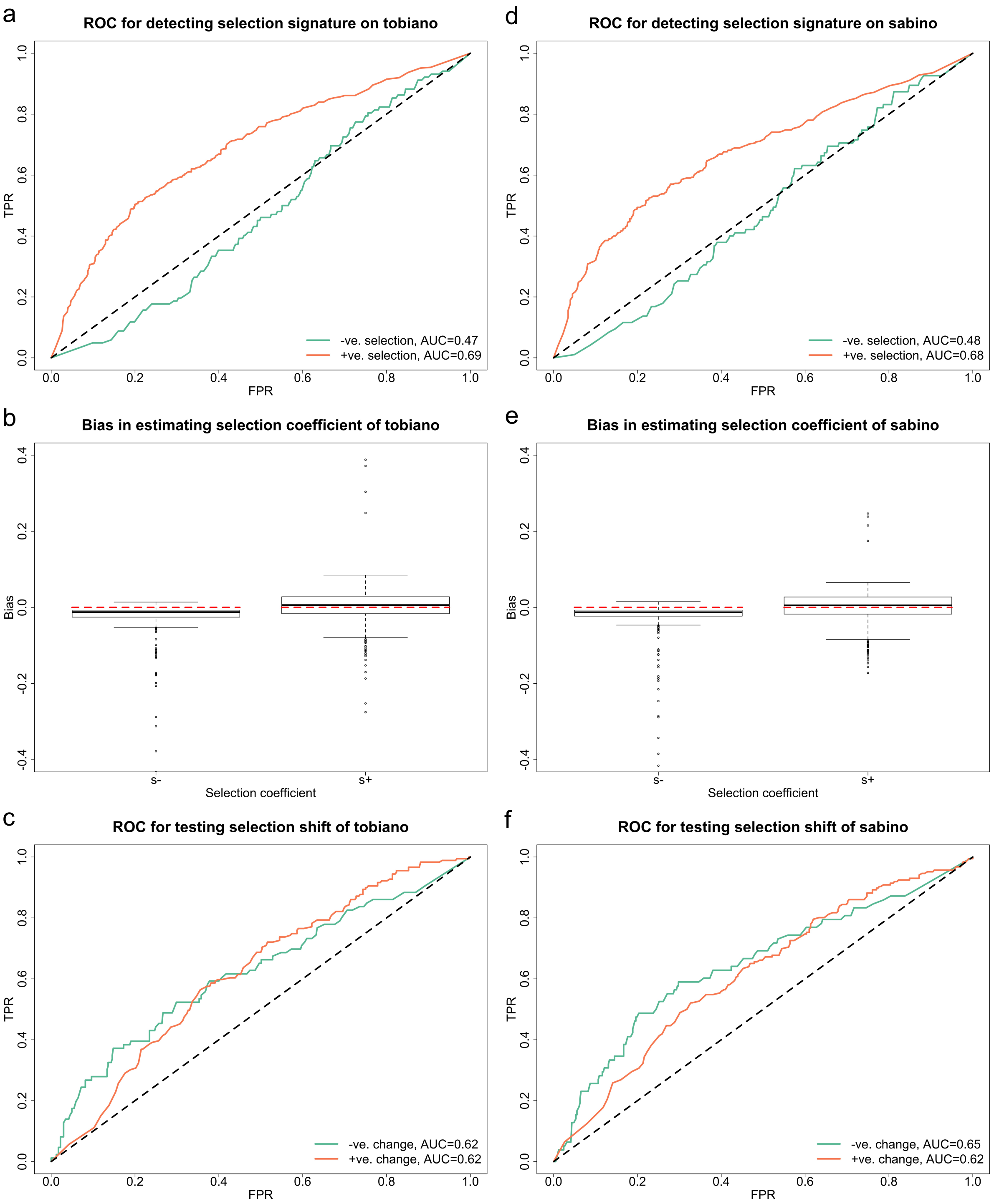


Figure S13: Empirical distributions for the bias in MMSE estimates of the selection coefficients for horse pinto coat patterns with ROC curves and AUC values for detecting selection signatures and testing selection changes, (a)-(c) for the tobiano coat and (d)-(f) for the sabino coat, respectively. To aid visualisation, we remove the replicates in which the absolute value of bias is larger than 0.4 from (b) and (e), *i.e.*, 83 replicates from (b) and 90 replicates from (e), respectively.
